# Long-lasting and responsive DNA/enzyme-based programs in serum-supplemented extracellular media

**DOI:** 10.1101/2021.11.22.469272

**Authors:** Jean-Christophe Galas, André Estevez-Torres, Marc Van Der Hofstadt

## Abstract

DNA molecular programs are emerging as promising pharmaceutical approaches due to their versatility for biomolecular sensing and actuation. However, the implementation of DNA programs has been mainly limited to serum-deprived in vitro assays due to the fast deterioration of the DNA reaction networks by the nucleases present in the serum. Here, we show that DNA/enzyme programs are functional in serum for 24h but are latter disrupted by nucleases that give rise to parasitic amplification. To overcome this, we implement 3-letter code networks that suppress autocatalytic parasites while still conserving the functionality of DNA/enzyme programs for at least 3 days in the presence of 10% serum. In addition, we define a new buffer that further increases the biocompatibility and conserves responsiveness to changes in molecular composition across time. Finally, we demonstrate how serum-supplemented extracellular DNA molecular programs remain responsive to molecular inputs in the presence of living cells, having responses 6-fold faster than cellular division rate and are sustainable for at least 3 cellular divisions. This demonstrates the possibility of implementing *in situ* biomolecular characterization tools for serum-demanding *in vitro* models. We foresee that the coupling of chemical reactivity to our DNA programs by aptamers or oligonucleotide conjugations will allow the implementation of extracellular synthetic biology tools, which will offer new biomolecular pharmaceutical approaches and the emergence of complex and autonomous *in vitro* models.

---

In the last few decades, the programmability and reactivity of DNA has positioned the DNA nanotechnology field as a promising avenue for the development of biomolecular pharmaceutical approaches.^1^ In particular, DNA molecular programming tools have been extensively used to create bioactuation systems for the delivery of cargoes^2^ and for the modification of cellular composition,^3^ as well as biosensing tools for cell sorting^4^ and molecular detection.^5^ For example, the specific amplification of nucleic acid sequences by polymerase-based DNA programs allows the detection of microRNA biomarkers down to attomolar concentrations with dynamic ranges up to 10 orders of magnitude.^5^ In addition, the amplification of DNA also grants the possibility of implementing DNA molecular computations, which has already been nicely exploited to diagnose cancer profiles from human samples.^6^ However, very few molecular programs work in direct contact with living cells.^1^ In contrast, current implementations favour the analysis of liquid biopsies within solutions that are not compatible with cellular growth^7^ or else use compartments that separate the program and the living system.^8–10^ As a result, these approaches limit the implementation of DNA pharmaceutical tools for *in vitro* studies.

In an effort to surpass this limitation, we recently demonstrated the embedding of a DNA/enzyme molecular program within the extracellular medium of an *in vitro* cell culture.^3^ Notably, we showed how the programmable extracellular medium was capable of guiding cellular composition across time and space, opening the pathway towards the development of extracellular synthetic biology tools. Nevertheless, the absence of serum reduces the biological significance of that study, since the myriad of bioactive substances provided by the animal serum has become an essential component for successful *in vitro* cell culture.^11^For this reason, the DNA nanotechnology field has recently focused on the stabilization of DNA nanostructures^12^ and DNA circuits within serum-supplemented media.^13^ However, to date, only Fern *et al.* have reported a solution for extending the functional life of DNA-only programs in the presence of serum components.^14^ The main limitation emerges from the unwanted interactions of the serum components that alter or degrade the DNA molecules, which leads to the rapid loss of the designed networks. In particular, this low resilience has been mainly attributed to the degradation of the DNA by the presence of nucleases in the serum, which restraints any potential for long-term application of DNA programs.

To overcome the degradation of the DNA by the serum, two major approaches have been developed. Firstly, the nuclease activity can be impaired by using DNAse inhibitors (such as actin) or by heat inactivating the serum.^15^ Secondly, efforts have been focused on the protection of the synthetic DNA strands by the introduction of structural changes, either chemical^16, 17^ or morphological.^14^ While the first approach is incompatible with nuclease-assisted DNA programs and lacks biological significance (as the heat inactivation denatures all proteins present in the serum), the second is not suitable for polymerase-based DNA programs as the *de novo* synthesized DNA cannot be protected *in situ*. For these reasons, to the best of our knowledge, long-lasting DNA/enzyme molecular programs have not yet been described in the presence of animal serum, limiting the potential of using DNA molecular programs in the presence of living cells for in situ biosensing and bioactuation.

Here, we demonstrated that a DNA/enzyme-based molecular program is functional in the presence of animal serum and living cells for at least three days. Firstly, we show that the existence of nucleases in the serum disrupt the polymerase-based DNA programs, since the emergence of non-programmed parasitic amplification is enhanced and overtakes the DNA program. To circumvent this, we restrained the undesired activity of serum nucleases by avoiding the creation of *de novo* double stranded DNA (dsDNA) through the use of 3-letter code templates. The reformulated DNA programs were capable of responding to sequence-specific single stranded DNA (ssDNA) and triggering the *in situ* production of ssDNA up to 1 *μ*M for at least 49 h in the presence of 10% fetal bovine serum. Secondly, we show that such serum conditions are needed to conserve the phenotypic behaviour of *in vitro* human embryonic kidney cells. Finally, we demonstrate that serum-supplemented DNA/enzyme-based extracellular programs remain responsive in the presence of living cells, paving the way for the development of DNA-regulated extracellular synthetic biology systems that would complement traditional intracellular approaches to create complex and autonomous *in vitro* models.

## Results

### Serum disrupts the functionality of DNA/enzyme molecular programs

To asses the functionality and robustness of DNA/enzyme-based programs in serum, we performed an exponential amplification reaction (EXPAR) in the presence of fetal bovine serum (FBS). In particular, we focused on the polymerase, nickase, exonuclease dynamic network assembly toolbox (PEN DNA toolbox)^18^ as it allows the assembly of elementary reactions (Figure S8) into complex functional reaction networks.^19, 20^ Figure 1a exemplifies the behaviour of a PEN DNA autocatalyst, where ssDNA **A_1_** is amplified exponentially in the presence of template **T_1_** and the three enzymes cited above. The fluorescence of the intercalator EvaGreen allows to follow, over time, the total concentration of double stranded DNA (dsDNA) in solution. This PEN amplification reaction is characterized by a sigmoidal curve, where an initial exponential phase is followed by a linear phase before reaching saturation. At saturation, the EvaGreen fluorescence reaches a plateau corresponding to a steady state of dsDNA concentration and due to the constant production and degradation of *de novo* DNA by the polymerase and exonuclease, respectively.^18^ In a first series of experiments, we tested the autocatalytic template **T_1_** in a buffer compatible both with PEN reactions and mammalian cell culture^3^ (named *Kin* buffer due to its high enzyme kinetics, see SI Section 3), for increasing concentrations of serum. In all conditions, we observed a decrease in the fluorescence intensity during the first 500 min (Figure 1b), reduction we account to an artifact due to the presence of 0.5x cell culture medium (Dulbecco's modified Eagle's medium, DMEM) in the buffer. However, the onset of the exponential amplification by the autocatalytic **T_1_** template can still be clearly distinguished by the sigmoidal curve starting after 500 min in the absence of FBS. Upon the introduction of 2.5% FBS, the DNA amplification dynamics were slowed down (loss in steepness of the linear region) and the dsDNA steady state regime was not persistent, since it was followed by a nonlinear signal increase after 1800 min. We attribute both these deficiencies to the emergence of untemplated replication of DNA,^21^ which gives rise to parasitic amplification networks that hijack the enzymes and energy source. As FBS concentration was further increased, the linear regime shortened and the nonlinear signal increase started earlier. At 10% FBS (standard cell culture concentrations) the initial exponential amplification regime of the synthetic **T_1_** template cannot be distinguished from the parasitic non-linear amplification. Denaturating polyacrylamide gel (PAGE) clearly demonstrated the production of copious DNA strands not related to the original amplification network (SI Section 2), and consequently the loss of the functionality of the DNA/enzyme molecular program.^22^

**Figure 1:**
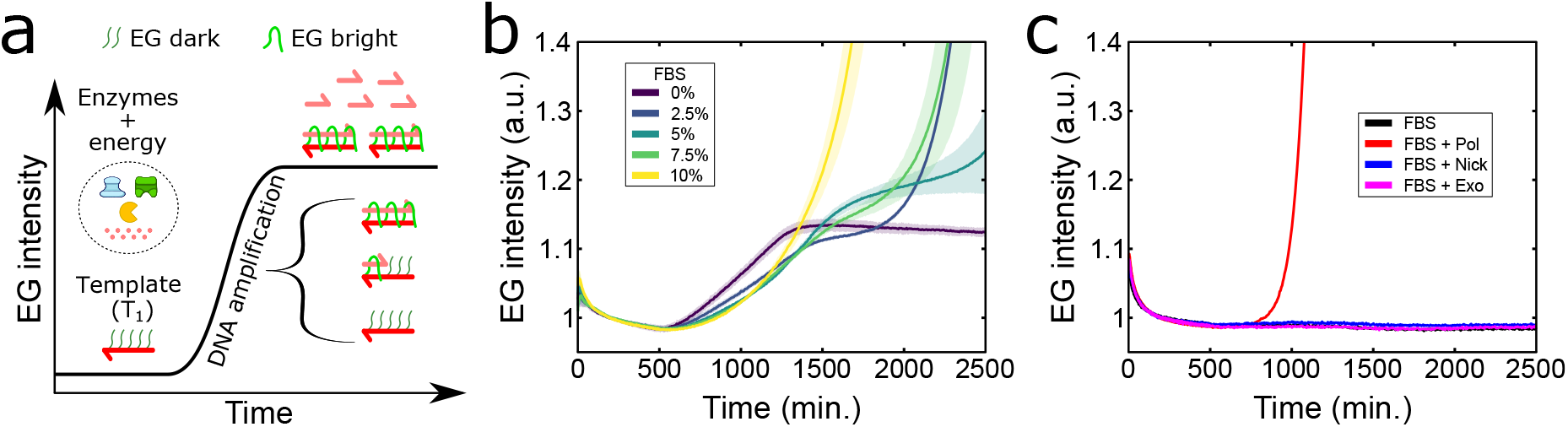
Serum promotes the emergence of parasitic amplification in the presence of polymerase-based DNA programs. (a) Scheme of the PEN DNA exponential amplification reaction depicting the nature and behaviour of the EvaGreen (EG) reporter, a fluorescent dsDNA intercalator, on the autocatalytic **T_1_** template. Harpoon-ended arrows represent single stranded DNA (ssDNA). (b) EvaGreen fluorescence *versus* time for the **T_1_** autocatalytic network at 37 °C in a concentration range of FBS. (c) EG fluorescence *versus* time for the incubation of 10% FBS with one of the three enzymes present in PEN reactions: polymerase (pol), Nb.BsmI nickase (nick) or exonuclease (exo). The shades in panel b correspond to one standard deviation of a triplicate experiment. Conditions panel b and c: *Kin* buffer with 0.1 mM dithiothreitol (DTT), in addition to 200 nM of autocatalytic **T_1_** template for panel b.

Although the exact mechanism of parasitic amplification is still not well understood, it is known that it emerges from the *de novo* synthesis of DNA by polymerases^23^ and by the presence of endonucleases that create tandem repeats^24^ and quasi-palindromic sequences,^21^ both containing the endonuclease recognition site. For instance, the polymerase and the nicking enzyme of the PEN reactions generate autocatalytic parasites in the presence of dNTPs.^22^ To test if the observed parasites in the presence of serum (Figure 1b) were due to the PEN nicking enzyme or to an endonuclease present in the serum, we performed the following experiment. In a controlled experiment, we generated a PEN parasite in the presence of Nb.BsmI (the PEN nicking enzyme), and a second with the addition of 10% serum. After parasite had emerged, we incubated both conditions in the presence of BsmI, the corresponding restriction enzyme (SI Section 2). We observed that after 3.5 hours of BsmI incubation, the DNA smear characteristic of PEN parasites observed in denaturating PAGE had been strongly reduced (Figure S1).^21^ In contrast, when we tested the degradability of parasites that had emerged in the presence of 10% FBS, we observed that, after 8 hours of BsmI incubation, the smear had only partially disappeared (Figure S2). We further tested the potential of FBS to give rise *per se* or aided by PEN enzymes to the creation of parasite. We incubated 10% FBS in the absence or in the presence of one of the three enzymes present in PEN reactions and followed EvaGreen fluorescence for exponential amplification of *ab initio* dsDNA (Figure 1c). Results demonstrated that while 10% FBS was not capable of giving rise to parasite autonomously, parasite emerged only when polymerase was added to the FBS solution. In addition, we observed that the parasitic emergence time was inversely proportional to the FBS concentration (Figure S9). We infer that the FBS has endonuclease enzymes that, together with the polymerase and dNTPS used on PEN reactions, generates autocatalytic parasites that disrupts the DNA/enzyme programs.

### Impairing serum parasite emergence by using a 3-letter code

We have recently demonstrated that the emergence of PEN parasites can be overcome by selecting a PEN enzyme whose recognition site only bears 3 of the 4 nucleotides per strand, coupled to a 3-letter code template.^22^ As a result, in the absence of the fourth dNTP in the solution, *de novo* DNA synthesis by the polymerase cannot create *in situ* dsDNA that may be cleaved by the nicking enzyme, thus yielding a parasite (Figure S10).^21, 24^ Thus, by designing a PEN autocatalytic network with DNA templates containing only adenine, guanine and cytosine in their sequence, and removing adenosine triphosphate from the dNTPs solution (Figure 2a), one obtains a functional DNA circuit while avoiding the emergence of unwanted parasitic sequences.

**Figure 2:**
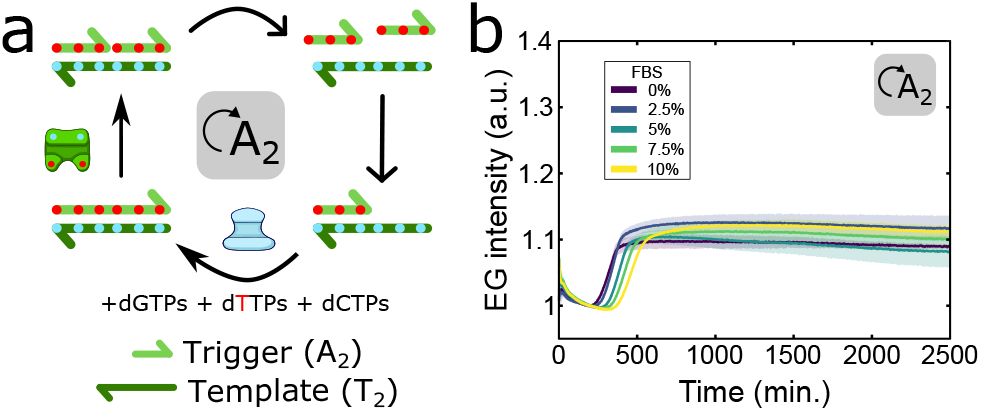
3-letter code DNA networks preclude the emergence of autocatalytic parasites originating from endonucleases present in the FBS. (a) Scheme of the 3-letter code autocatalyst resistant to parasitic emergence in FBS due to the absence of dATPs in the solution. In this network, the dATPs (blue spots) are restricted to the synthetic **T_2_** templates and the dTTPs (red spots) to the *de novo* **A_2_** ssDNA synthesis. Harpoon-ended arrows represent ssDNAs. Irreversible and reversible reactions are indicated by solid and empty arrowheads, respectively. (b) EG fluorescence *versus* time for the **T_2_** autocatalytic network with the Nb.BssSI nickase at 37 °C in a concentration range of FBS in the absence of dATPs. Note the absence of parasites compared to Figure 1b. The shades in panel b correspond to one standard deviation of a triplicate experiment. Conditions: *Kin* buffer with 0.1 mM DTT.

To investigate if a 3-letter code approach can also impair the emergence of serum-promoted parasites, we first designed an autocatalytic template (**T_2_**) that was based on the Nb.BssSI nickase, as all thymine bases are located in the same strand of the recognition site, and that worked at 37 °C. Figure 2b shows the amplification dynamics of template **T_2_** in the presence of 3 dNTPs and increasing concentrations of serum. The fluorescence intensity displayed a sigmoidal curve characteristic of PEN exponential amplification. Importantly, and contrary to 4-letter code templates (Figure 1b), the amplification dynamics were not significantly affected, for up to 42 h, even in the presence of 10% FBS. We do observe a modest delay on the onset of the exponential amplification upon the addition of 10% FBS associated to a slower nickase kinetics (Figure S3). As expected, parasite emergence was still observed in the presence of 4 dNTPs (Figure S11).

### PEN molecular programs remain responsive for at least 45 h

*In vitro* cellular doubling time largely depends on cellular type and growth conditions, ranging from 10 h to few days.^25^ For example, the doubling time for human cervix epithelial carcinoma cells (HeLa cell line) in our conditions was ∼17.5 h (Figure S12). In order to develop DNA extracellular programs that may interact with cells, one needs a DNA program that computes faster than cellular growth (*i.e.* <15 h) and that remains responsive for at least two cell cycles (*i.e.* >40 h). To introduce a fast and responsive behaviour into our DNA program, we took advantage of a repression module to implement a controllable bistable switch (Figure 3a).^18^ In this network, we define the *ON* state as the result from the sustained exponential amplification of **A_2_** by the autocatalytic reaction of template **T_2_**, as it has been shown above. Contrary, in the *OFF* state, the presence of high concentrations of the repressor strand (**R_2_**) suppresses this autocatalytic reaction due to the combined elimination of **A_2_** by the exonuclease and its conversion to waste (W) by **R_2_**. However, the *OFF* state can be reverted to the *ON* state by the addition of a DNA activator 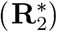, complementary to R2, that reactivates the exponential amplification of **A_2_**.^3^ The program is thus responsive to the molecular stimulus of **R_2_**. Note that, due to the low dNTPs consumption rate of the system in the *OFF* state, we expect the DNA program to remain responsive for long periods of time.

**Figure 3:**
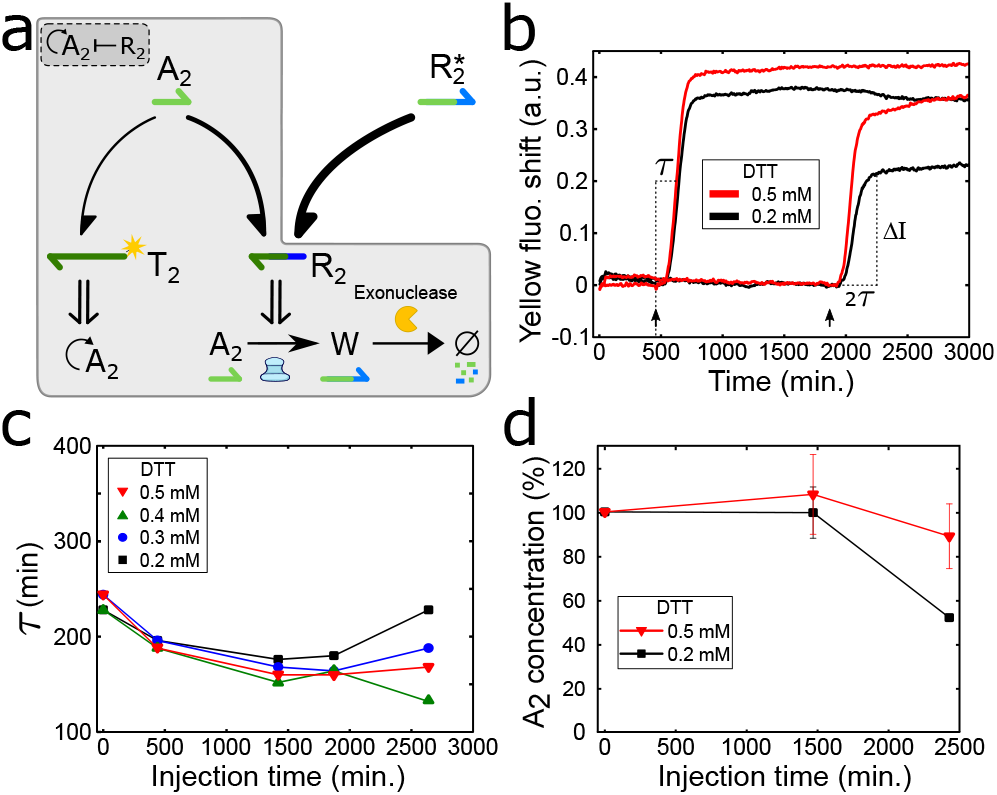
Long-lasting and responsive DNA programs in the presence of 10% FBS. (a) Scheme of the responsive DNA program. The combination of an autocatalytic node (**T_2_**) with a repressor node (**R_2_**) allows the creation of a bistable switch (grey enclosure), where the higher affinity of **A_2_** with **R_2_** compared with **T_2_** causes the extension of **A_2_** to a non functional ssDNA (waste **W**) and its degradation by the exonuclease. The addition of 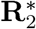 that binds to **R_2_** reduces the free concentration of the latter, promoting the exponential amplification of **A_2_** by **T_2_**. The thickness of half arrow-headed lines indicates DNA affinities. (b) Fluorescence shift from the fluorescently-labeled **T_2_** *versus* time for the responsive program triggered at different times with 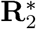 and for different DTTJ on the *Kin* buffer. The amplification onset time, *τ*, and the fluorescent amplitude of the response at dsDNA steady state, Δ*I*, are defined on the graph (see methods). (c) *τ versus* the time at which 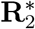 was introduced into the solution (injection time) at different DTTJ. (d) **A_2_** relative concentrations at steady state with respect to the value obtained for an injection time at t = 0 h, for different injection times and DTTJ (Figure S14). Data in panel c determined from panel b and Figure S6. Solid lines in panel c and d are guides to the eye. Error bars in panel d correspond to the standard deviation of a triplicate experiment. Conditions: [**R_2_**]_0_ = 100 nM. 200 nM of 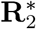 was added at the injection times (arrowheads in panel b).

To determine if the DNA program remained responsive in the presence of 10% FBS, we spiked the *OFF* state with 200 nM of 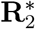 at different time points (Figure 3b and Figure S6). Since the dithiothreitol (DTT) prevents the oxidation of the enzymes,^26^ we hypothesized that increasing its concentration could help keeping the amplification onset time (*τ*) and the fluorescent amplitude of the response (Δ*I*) constant at longer times. We observed that the responsive time for up to 1868 min changed little for DTT concentrations ranging 0.2-0.5 mM (Figure 3c). In contrast, at 2700 min, increasing DTT concentration from 0.2 up to 0.5 mM made the response 26 % faster. Nevertheless, in all cases, the system was capable of responding at least 4.6-fold faster than HeLa cellular division rate and for at least 2.5 cell cycles.

The amplitude of the response (the steady-state signal of the fluorescently-labeled template, see SI Section 3) is also important for bioactuation purposes. Since we noticed that the Δ*I* at twice the onset time (2*τ*) was decreasing with the injection time (Figure 3b and Figure S13), we evaluated the DNA concentration behaviour across time. To do so, we quantified the **A_2_** ssDNA available after a 49 h run for an experiment spiked at 0, 24 and 40 h (Figure S14). When spiked at t = 0 h, the DNA program produced respectively 0.8 *μ*M and 1.1 *μ*M of **A_2_** in the presence of 0.2 and 0.5 mM DTT, which is ∼5-fold greater than the template concentration (Figure 3d). We observed that these levels were similarly reached for experiments spiked after 24 h. Regarding 40 h spiked experiments, the production of **A_2_** could only reach 53 ± 1% when 0.2 mM DTT was used. However, this production increased up to 89 ± 15% in the presence of 0.5 mM DTT, showing the great robustness of PEN networks in the presence of serum. Furthermore, since the onset of the exponential amplification is dependent on 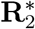 concentration (Figure S15), the designed DNA program will also be capable of quantifying changes in molecular composition.

### Increasing the biocompatibility of the buffer

We have demonstrated that greater DTT concentrations improve the responsiveness of the DNA program in the presence of serum. However, since high levels of [DTT] decrease cell viability, as they transiently activate endoplasmic reticulum stress^27^ and cause cell detachment,^3^ we decided to further increase the biocompatibility of our buffer to mitigate cytotoxicity (although the introduction of FBS already partially attenuates the adverse effect of DTT, Figure S16). To do so, we increased the concentration of the cell culture medium (from 0.5x to 0.89x), we removed non-critical PEN buffer components and lowered the concentration of magnesium (SI Section 3) to create a new buffer that we named *Cell +* buffer. Interestingly, we observed that the chosen 3-letter code nickase (Nb.BssSI) was more robust to buffer modifications than the traditional Nb.BsmI nickase used for PEN reactions and that they were also functional in RPMI-1640, another standard cell culture media (Figure S7).

In contrast with the results shown above in the *Kin* buffer, we observed that the *τ* and **A_2_** concentrations at steady state in the *Cell +* buffer were strongly dependent on [DTT] and injection time (Figure 4 and Figure S13). In particular, we observed that at low DTT concentrations (0.2 mM) *τ* values increased by 139% when the DNA program was activated after 45 h. However, at higher concentrations (>0.3 mM) the DTT concentration had no further effect. Quantification of the available **A_2_** at steady state revealed that it decreased steadily at longer injection times. When the injection occurred at 24 h, **A_2_** was produced at ∼75% of the steady state level when injected at t = 0 h. Regarding the injection at 40 h, **A_2_** dropped to 26 ± 5% and 60 ± 6% for the 0.2 and 0.3 mM of DTT, respectively. As the response (*τ* and [**A_2_**]) is not largely affected when the [DTT] is increased above 0.3 mM, we decided to use this concentration of DTT to have a compromise between functionality and biocompatibility for the new *Cell +* buffer.

**Figure 4:**
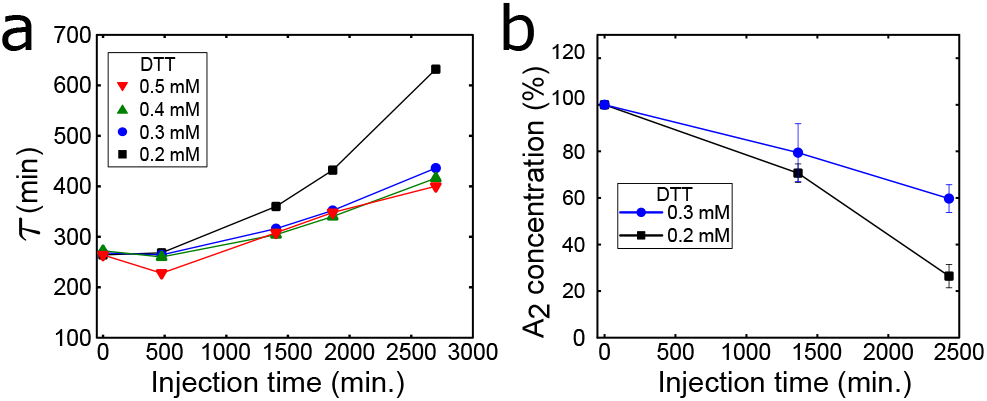
Responsiveness of the DNA program in the *Cell +* buffer. (a) *τ versus* injection time of 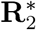 in *Cell +* buffer in a range of [DTT]. (b) **A_2_** concentration for the injection of 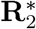 after 24 h and 40 h quantified from Figure S14. Data in panel a determined from Figure S6. Solid lines are guides to the eye. Error bars in panel b correspond to the standard deviation of a triplicate experiment. Conditions: [**R_2_**]_0_ = 100 nM. 200 nM of 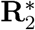 was added at the injection times.

Once that we have evaluated the performances of the DNA/enzyme program in the serum-supplemented *Kin* and *Cell +* buffers, it is now necessary to measure cellular viability in these buffers. To do so, HeLa cells were stained with propidium iodide and their fluorescence evaluated using flow cytometry at different culture times (Figure 5a and Figure S12). As expected, in the control condition the number of living cells increased with time. However, we noticed a 1.6-fold reduction on cellular growth rate after 48h that didn’t occur in the absence of FBS (Figure S12), most likely indicating an impairment on cell growth due to the exposition to serum-supplemented media for long periods of time (e.g. high metabolism). To test the biocompatibility of the *Kin* buffer, we decided to use 0.5 mM DTT to conserve the fast and long-lasting responsiveness of the DNA program assessed in the previous section. Results revealed a significant reduction on cellular viability (down to 71%) and 3.6-fold lower cell number after 24h compared to the control, which we attribute to the toxicity and cellular detachment introduced by the high DTT concentration. On the other hand, the *Cell +* buffer at 0.3 mM DTT conserved high viability (above 93%) and similar cell number after 24h of incubation as the control. Although we found an average ∼2-fold reduction in growth rate compared to the control, a ∼2-fold increase in viable cell number after 48 h was achieved compared to previously reported experimental conditions in the absence of FBS (Kin buffer).^3^

**Figure 5:**
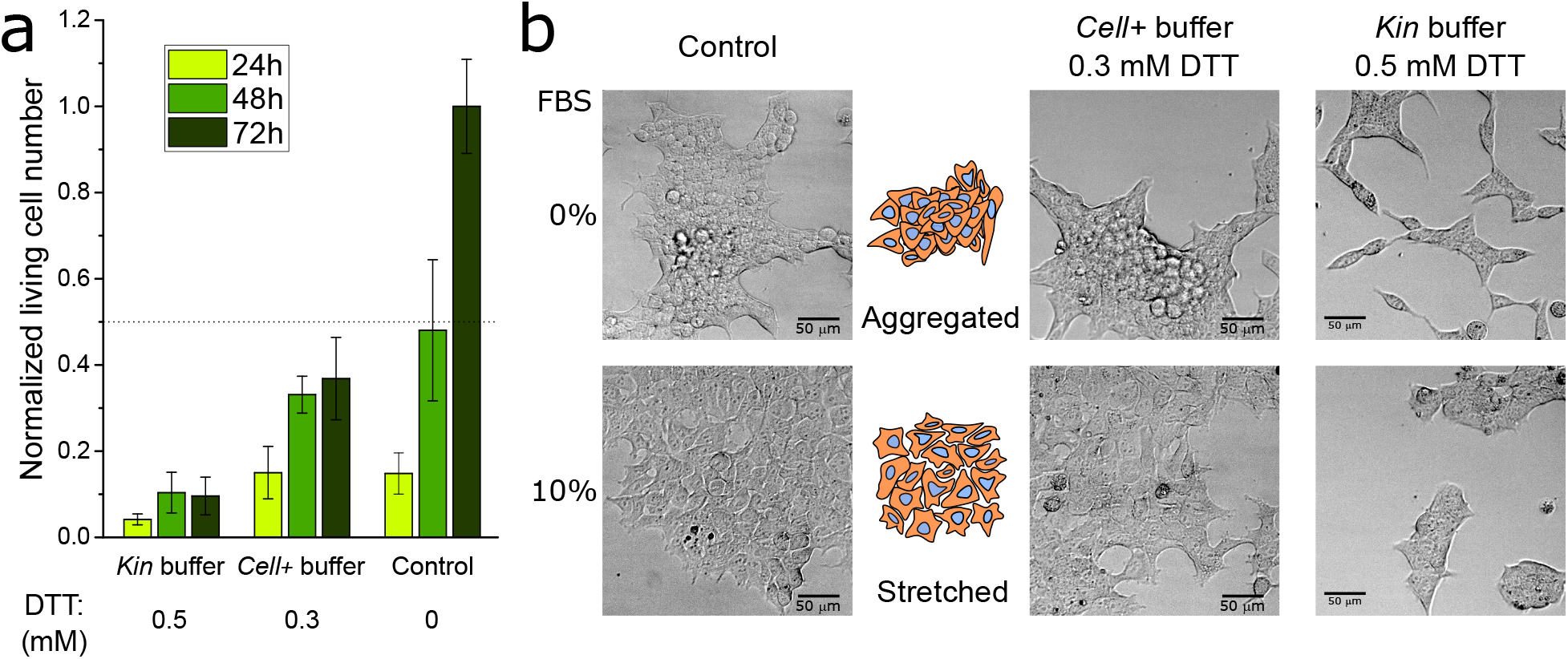
Evaluation of cell viability for the three buffers used in this study. (a) Normalized living cell number of HeLa cells determined by cellular staining with propidium iodide and flow cytometry for different incubation times and conditions in the presence of 10% FBS. Data determined from Figure S12. (b) Bright-field images demonstrating the morphology of HEK cells after 72 h of incubation in the absence or the presence of 10% FBS for the control growth medium, *Cell +* buffer and *Kin* buffer. Images obtained from the time-lapse of Figure S19.

To stress the importance of developing serum-compatible DNA/enzyme programs, we studied the growth of human embryonic kidney 293 cells (HEK 293 cell line), as their behaviour (e.g. signalling pathways) is significantly affected by serum starvation.^28^ We observed that in the absence of serum, HEK cells aggregated in standard cell culture medium, phenotype that was not appreciable when supplemented with 10% FBS (Figure 5b). We noted that the stretched morphology was still conserved for at least 48 h when the FBS was reduced down to 2.5% (Figure S17), opening the possibility to reduce the FBS concentration without perturbing cellular phenotype. When we evaluated the morphology and viability of HEK cells in the two DNA buffers, we noticed that, contrary to HeLa cells, HEK cells are less resilient to the presence of DTT (Figure S18), most likely due to their lower adhesion to surfaces. Nevertheless, HEK cells conserved high cellular viability and stretched morphology in the serum-supplemented *Cell +* buffer in contrast to what happened in the *Kin* buffer (Figure 5b and Figure S12).

### Serum-supplemented DNA programs are functional in the presence of cells

To verify that the DNA/enzyme program was still functional in the presence of living cells and 10% FBS, we first tested the bistable switch. To do so, we seeded the cells in a cell culture multiwell plate, and allowed them to adhere for 24 h before introducing the *Cell +* buffer containing the DNA program (Figure 6a). When the DNA program was set to be in the *ON* state ([**R_2_**]_0_ = 0 nM), the exponential amplification of DNA occurred within 2 h and reached a steady state after 23 h (Figure 6b). When the program started in the *OFF* state ([**R_2_**]_0_ = 150 nM), the amplification of DNA was suppressed for at least 71 h, showing the robustness of the *OFF* state in the presence of living cells. Quantification of the available **A_2_** at steady state revealed that the same concentration of ssDNA was produced for the *ON* state in the absence (Figure 3d) and in the presence of cells (Figure 6c), and that no **A_2_** was produced in the *OFF* state.

**Figure 6:**
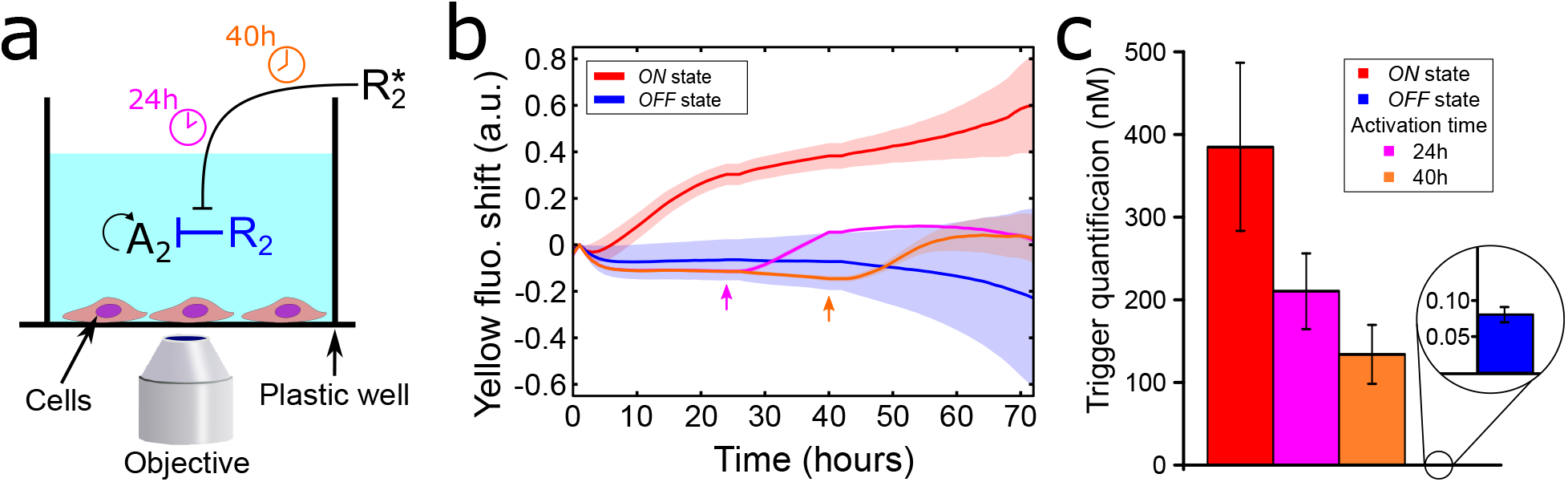
Long-lasting and responsive behaviours of serum-supplemented extracellular DNA programs in the presence of living cells. (a) Cartoon of the experimental setup (not to scale). The cells are cultured in the extracellular DNA program where the **A_2_** autocatalyst is not suppressed (*ON* state) or permanently repressed (*OFF* state). In the later case, the *OFF* state can be reverted by the external addition of DNA activator 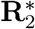 after 24 h (pink) or 40 h (orange). (b) Fluorescence shift of **T_2_** *versus* time showing the production dynamics of **A_2_** in the *Cell +* buffer in the presence of 10% FBS and HeLa cells. The curves show the unsuppressed *ON* state (red) and the repressed OFF state (blue) and its responsiveness by the addition of 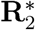 at 24 h (pink) and 40 h (orange). Arrowheads indicate the addition time of DNA activator 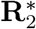. (c) **A_2_** trigger concentrations after 71h of incubation of the DNA/enzyme-based molecular program in the presence of cells and 10% FBS. Data in panel c determined from Figure S20. Conditions: The ON and OFF states started with [**R_2_**]_0_ = 0 nM and 150 nM, respectively. The DNA activator 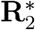 was introduced at 300 nM.

To test the responsiveness of the DNA program in the presence of living cells, the *OFF* state was turned *ON* by the addition of 300 nM of 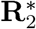 after 24 h or 40 h. In both cases, we observed that the amplification of DNA was initiated within 3 h, which is ∼6-fold faster than cellular growth rate, and reached a steady state after ∼15 h, where it remained until the end of the experiment. However, we noticed a ∼50% reduction in the dsDNA steady-state fluorescence compared to the *ON* state. The quantification of **A_2_** showed that the injections at 24 and 40 h produced 210 and 134 nM of **A_2_**, respectively. This is 2-3 times less than what was produced in the *ON* state but still remarkable and largely sufficient to perform downstream molecular computations. We hypothesize that this reduction is due to the presence of cells, either due to the cell debris and secretions that may interfere with the DNA/enzyme program, or due to a greater DNA uptake by cells due to the presence of FBS.^29^ In addition, we contemplate that in early activated DNA programs, due to the high concentration of *de novo* **A_2_** ssDNA compared to the initial synthetic template **T_2_**, the core of the DNA program (*i.e.* the template) is significantly less affected by the presence of cells. Similar results were obtained for the lesser biocompatible *Kin* buffer, although with a faster response and higher **A_2_** concentrations at steady state (Figures S20 and S21). Finally, clock reactions controlling the onset time of the exponential amplification in a pre-encoded manner were also functional in the presence of serum (Figure S21). These results demonstrate the feasibility of programmable and responsive DNA/enzyme based molecular programs in the presence of cells and 10% serum.

## Conclusions

In this paper, we demonstrate the co-existence of responsive DNA/enzyme-based programs and living cells in the presence of 10% animal serum for at least 3 days, opening a route to the implementation of *in situ* molecular characterization tools. We have shown that the long-lasting programmability of polymerase-based DNA programs in the presence of serum is lost in under one day due to the unwanted nuclease activity present in FBS, which gives rise to parasitic amplification that hijack the enzymes and energy source of the PEN DNA reaction networks. To overcome the emergence of parasites, we used DNA circuits that have been designed to only have 3 deoxynucleotides per strand instead of the traditional 4-letter code. By removing autocatalytic parasites, we could generate a DNA/enzyme program that was responsive for >45h in serum-supplemented buffers. Results revealed that the responsive behaviour to the activation of a bistable DNA switch was capable of producing and maintaining in situ up to 1 *μ*M of ssDNA for at least 49 h. In addition, cellular viability results revealed a ∼2-fold increase in viable cell number of our serum-supplemented DNA programs in the new *Cell +* buffer in comparison to previous conditions in the absence of FBS.^3^ Importantly, we have demonstrated that serum-supplemented buffers are capable of conserving the normal stretched phenotype of HEK cells, stressing the importance of developing serum-compatible molecular programs when adventuring for *in vitro* cell culture experiments. Finally, our cell culture results corroborate that serum-supplemented extracellular DNA molecular programs in the presence of living cells are functional, are pre-programmable, can respond to extra-cellular perturbations faster than cellular division rates and are sustainable for at least 3 cellular divisions (71 h).

The DNA network and buffer optimization shown here can be further extended to other DNA programs and other cell types with adequate modifications. For instance, due to the lower abundance of 3-letter code nucleases,^30^ we hypothesize that the implementation of DNA-only networks based on 3 deoxynucleotides per strand can reduce the presence of restriction sites of serum nucleases that passively degrade the DNA circuits.^14^ Regarding other cell types, we have shown that the DNA program is capable of working in two of the most standard cell culture media (DMEM and RPMI-1640), opening the possibility of its implementation to other cell types with sufficient adaptation. In particular, we stress the fact that we have shown functionality of the DNA program under the conventional use of 10% FBS for cell culture, but not all cells require 10% FBS to conserve growth and phenotype. This grants the opportunity for reducing the concentration of FBS in the presence of the DNA program, which would imply the lower need of [DTT] for DNA responsiveness and hence the buffer would present even lower cytotoxicity for sensitive cell types (*e.g.* neurons).

With these outcomes, we now envision the implementation of extracellular DNA programs capable of responding to changes in the molecular composition in the presence of living cells. In particular, the extracellular DNA programs will allow the *in situ* biomolecular recognition during *in vitro* cell culture, avoiding *ex situ* non-biocompatible amplification mechanisms that lack temporal and spatial resolution.^5^ Furthermore, since the PEN DNA toolbox offers a large set of functional reaction networks, such as biochemical concentration patterns^20, 31, 32^ and trigger-driven networks^33, 34^ with out-of-equilibrium properties, DNA molecular programs have potential to be advantageously used to create extracellular synthetic biology approaches. These approaches can be coupled to the abundant array of reactivity offered by oligonucleotides, either chemical^35^ or structural,^36^ to offer new biomolecular pharmaceutical tools. Likewise, communication pathways from the cells to the synthetic programs may be implemented by means of natural precursors (digestion, internalization) or synthetic approaches relying on biomolecular triggers,^37^ which could be merged to the reactivity of oligonucleotides for the creation of complex and autonomous *in vitro* models.

## Methods

The design of all DNA strands (Table S2) was done heuristically and assisted by Nupack,^38^ and purchased from Integrated DNA Technologies, Inc (U.S.) or Biomers (Germany). Both nickases (Nb.BssSI and Nb.BsmI) and the Bst DNA polymerase large fragment were purchased from New England Biolabs. The *Thermus thermophilus* RecJ exonuclease was produced in the lab following previous protocols.^39^ Standard enzymatic concentrations used in this study were 8 U/mL polymerase, 100 U/mL Nb.BsmI and 31.25 nM exonuclease for Nb.BsmI experiments, and 6.4 U/mL polymerase, 20 U/mL Nb.BssSI and 50 nM exonuclease for Nb.BssSI experiments.

The cell growth medium contained Dulbecco’s modified Eagle’s medium (DMEM F12, PAN Biotech P04-41150) supplemented with 1% Penicillin-Streptomycin. While *Kin* buffer was the previously developed biocompatible medium,^3^ *Cell +* buffer was a further optimization to reduce toxicity without drastically losing DNA programmability (SI Section 3). Only *Kin* buffer and *Cell +* buffer contained dX TPs (where *X* could be 3 or N, see Table S1) at mM. Unless otherwise mentioned, all experiments were performed at 37 °C and 200 nM DNA Template. DNA sequences, buffer composition and further experimental procedures can be found in the Supplementary Information.

### PEN reactions in the absence of cells

Experiments were done in 20 *μ*L solutions and the dynamics of the PEN reactions were exposed by fluorescent changes recorded by a Qiagen Rotor-Gene qPCR machine or a CFX96 Touch Real-Time PCR Detection System (Bio-Rad). EvaGreen was used at 0.5x to detect parasite emergence. To remove artifacts due to the perturbation of the solution when the DNA activator 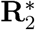 was introduced and the opening of the thermal cycler machine, a home made Matlab (The Mathworks) script was used to mathematically equal the fluorescence before and after the perturbation. To calculate the fluorescence shift, the raw fluorescence intensity was corrected by an early time point and subtracted from 1, as done previously.^20^ Both procedures are detailed in SI Section 1.2. For EvaGreen intensity values, the raw fluorescence intensity was only corrected by an early time point. To calculate the onset amplification time, *τ*, the sigmoidal curve was fitted with a polynomial fit, followed by the derivative of the polynomial fit. The time point with the highest derivative value was chosen as the *τ* value. The amplitude of the response (Δ*I*) was measured at the time point when the exponential amplification reached dsDNA steady state. We defined this Δ*I* time point as the time point when twice the onset amplification time from the perturbation of the system has been attained.

### Cell culture handling and experiments

Human cervix epitheloid carcinoma cells (HeLa cell line) and Human embryonic kidney 293 cells (HEK 293 cell line) were grown at 37 °C and 5% CO_2_. To reduce the cellular shock upon removal of FBS, the cells were grown in two sequential steps of FBS (Dominique Dutscher: S1810-500) (10% and 5%) before maintaining cell culture growth at 2.5% FBS. Bright-field images for observing cellular morphology were obtained with a Zeiss Axio Observer Z1 microscope with a 10X objective. When cells reached 80-90% confluence, they were detached with trypsin-EDTA (PAN Biotech: P10-019100) and diluted into fresh 2.5% FBS cell growth medium. For experiments, 800 cells were seeded in 384 cell well plates (ThermoFisher: 142762) and allowed to adhere onto the surface for 24 h before replacing the medium with 50 *μ*L of the experimental condition.

To quantify cellular viability by fluorescence activated cell sorting (FACS), the experimental condition was replaced with 50 *μ*L trypsin-EDTA (stock solution) and incubated for 8 minutes prior inactivation with 50 *μ*L of 10% FBS-supplemented cell growth medium. The cell suspension was mixed with 150 *μ*L FACSFlow (Fisherscientific: 12756528) and 0.5 *μ*L of propidium iodide (∼15 mM, ThermoFisher: P3566). The propidium iodide was excited with a 488 nm argon ion laser equipped within the Becton-Dickinson flow cytometer (FACSCalibur), and the fluorescent emission recorded within the fluorescence channel FL-2 (band pass 585/42 nm). Cells were quantified for 3 min at a flow of 60 *μ*L/min, and data was treated and analysed with a home-made Matlab routine. Cell count number was divided by the control growth medium to obtain the normalized living cell number.

### PEN reactions in the presence of cells

Cells and PEN reactions were monitored using a fully automated Zeiss Axio Observer Z1 epifluorescence microscope equipped with a ZEISS Colibri 7 LED light, YFP filter set, and a Hamamatsu ORCA-Flash4.0V3 inside a Zeiss incubation system to regulate temperature at 37 °C, in the presence of high humidity and at 5% CO_2_. Fluorescence images were recorded every 1 h with a 2.5X objective.

Images and data were treated as previously reported.^3^ Briefly, an ImageJ / Fiji (NIH) routine was implemented to stack the time-lapse images of each well. Subsequently, a Matlab routine was used to create an intensity profile across time of each well, which was divided by an initial time point to correct from inhomogeneous illumination between wells. To correct for time-dependent artifacts (e.g evaporation), the profiles were normalized with a negative control (absence of enzymes, *n* = 2). Lastly, fluorescent jumps and fluorescence shifts were calculated as described above.

### Available **A_2_** ssDNA quantification

Samples were extracted from the condition of interest and diluted down to 0.025% or 0.075% into a fresh isothermal amplification reaction containing [**T_2_**]_0_= 50 nM. The amplification onset times (*τ*) were plotted within a trigger titration calibration curve for extrapolating the quantification of the available **A_2_** ssDNA. To avoid perturbations from the FBS or the buffer sample, the fresh isothermal was performed with standard PEN DNA toolbox buffer that contains 3 mM DTT.^20^

## Supporting information

The Supporting Information contains further experimental methodology, a discussion on parasitic emergence by FBS and details on the buffers, DNA sequences, 23 supporting figures.

## Acknowledgements

We thank Stéphanie Bonneau and Ramón Eritja for supplying the HeLa cells, and Matthieu Morel the HEK 293 cells, Yannick Rondelez and Guillaume Ginés for insightful discussions, and Nelly Henry for the assistance with the FACS. We also thank the financial support from the European Research Council (ERC) under the European’s Union Horizon 2020 program (grant no. 770940, A.E.-T.), by the Ville de Paris Emergences program (Morphoart, A.E.-T.), by a Marie Sklodowska-Curie fellowship (grant no. 795580, M.vdH.) from the European Union’s Horizon 2020 program, and by a PRESTIGE grant (grant no. 609102, M.vdH.) from the European Union’s Seventh Framework Programme.

## Notes

### Competing Interest Statement

The authors have declared no competing interest.

