## supplementary information for "Long-lasting and responsive DNA/enzyme-based programs in serum-supplemented extracellular media"

### Contents


**1 Methods** **3**

**2 Parasite examination** **6**

**3 Higher robustness of Nb.BssSI to cellular media** **9**

**4 Table S1 to S2** **12**

**5 Figures S4 to S23** **14**

**References** **34**

### 1 Methods

#### 1.1 Determining the enzymatic rates of nickases

The nicking rates of Nb.BsmI and Nb.BssSI were measured in *Kin* buffer and *Cell+* buffer either in the absence or in the presence of 10% FBS at 37 °C. We used a reference substrate consisting of a DNA molecular beacon whose stem carried the nicking enzyme recognition site for Nb.BsmI ($ref_1$) and Nb.BssSI ($ref_2$). The nicking event caused the release of a short oligonucleotide containing the fluorescent dye, causing an increase in the fluorescence throughout the reaction.

The fluorescence signals of 20 $\mu$L solutions were tracked using a Qiagen Rotor-Gene qPCR. In the absence of FBS, the signal increased linearly until it reached a plateau, while in the presence of 10% FBS there was a background increase in fluorescence due to the presence of nucleases within the FBS (Figure S4). In the later case, we subtracted the background increase (in the absence of Nb.BsmI or Nb.BssSI) to the fluorescence signals before normalization between 0 and 1 and converting it to DNA concentrations by its multiplication by the reference substrate concentration (100 nM). A linear function of the treated data was fitted to the first tens of minutes and the gradient determined the enzymatic rates (Figure S4).

#### 1.2 Data treatment

A home made matlab script was implemented to remove fluorescent artifacts that occur due to temperature instabilities that alter the fluorescence intensity of fluorophores.[1] These temperature instabilities were mainly due to the opening of the experimental container (either PCR tube or cell culture multiwell plate) and temperature fluctuations in the room.

First, the raw data was imported and the time points at which the heating source (either the thermal cycler or the microscope) was opened were specified (Figure S5a). To introduce the DNA activator $\mathbf{R}_2^*$, the experimental container must be removed from the heating source to help handling and maintaining sterile conditions, which causes a drop in the temperature of the solution. In addition, although the added $\mathbf{R}_2^*$ volume was restrained to under 2.5% of the final volume, this injection also caused a reduction in the temperature of the solution and diluted the fluorophore concentration, both causing a reduction in the raw fluorescence intensity. To remove this thermal and dilution artifact, the fluorescent intensity after the opening of the heating source was mathematically equalized to the fluorescent intensity before the opening (Figure S5b). The number of time points that were mathematically equalized was dependent on the experimental procedure: a total of 6 time points (3 before and 3 after the injection time) were equalized for the thermal cycler experiments, while only 3 time points (1 before and 1 after) were equalized for microscopy experiments, as in the later case there was sufficient time for temperature stabilization (1 h time intervals). Note that even when no DNA activator is added, fluorescence intensity undergoes minor fluctuations that need to be corrected.

After removing time-specific artifacts, we observed an overall fluorescent artifact that was enhanced in experiments performed in the Qiagen Rotor-Gene qPCR machine. Since this qPCR machine is only capable of heating, the thermal fluctuations during the day (20 °C - 30 °C) cause an important effect on temperature stabilization, as the experimental temperature used in this work is low (37 °C). To these experiments (mainly MT Figures 3 and 4 and Figure S 6) we removed these long thermal fluctuations. To do so, we created a baseline curve that contained two components: since we observed that the major fluctuations were occurring for quenched fluorescent templates (*ON* state), we selected an experiment that had been activated at t= 0 h and had the highest DTT concentration (0.5 mM), to reduce any perturbations due to the presence of FBS. To avoid the autocatalytic regime of the curve (the quenching of the fluorophore), we only selected the curve from the 1000 minute time point to the end of the experiment. For the first 1000 minutes of the baseline curve, we created a mathematically flat line with the 1000 minute fluorescent intensity value. We then proceeded to divide all the curves by this reconstructed baseline curve to remove the long thermal fluctuations (Figure S5c). Note that for the experiment chosen to create the baseline curve there is no experimental significance in plotting the curve after the 1000 minute time point, reason why it has not been plotted in Figure S6.

To calculate the template fluorescence shift, the fluorescence intensity was firstly corrected by an early time point after its activation (Figure S5d). Since experiments done in the presence of cells were corrected from inhomogeneous illumination between wells at an early time point of the experiment, this step was not required. Subsequently, the corrected fluorescence was subtracted from 1 to obtain the fluorescence shift (Figure S5e).

#### 1.3 Polyacrylamide denaturing gel

Polyacrylamide denaturing gel electrophoresis at 20% was run for 2 h at 200 V in 0.5X TBE buffer, stained with 1000x Sybr Gold (ThermoFisher: S11494) for 10 min, and imaged using a Gel Doc™ EZ Gel Imager (Bio-Rad). Note that we use species $\mathbf{A}_1^1$ because upon the hydrolysis of a phosphodiester bond during the nicking event, the phosphate group remains in the 5' of the second trigger, since if the phosphate group remained on the 3' of the first trigger no autocatalytic behaviour would be attainable.

### 2 Parasite examination

To understand the emerging autocatalytic parasites in the presence of FBS, we first studied the degradation properties of the parasites emerging in conventional DNA buffers. To do so, we incubated 50 nM of template $\mathbf{T}_1$ with 100 U/mL Nb.BsmI nickase, and increased the temperature up to 44 °C and the polymerase concentration to 16 U/mL to enhance parasite apparition (Figure S1a). Under these experimental conditions, the programmable autocatalyst reached steady state after 75 min but was overrun by the parasite after 200 minutes. After a total incubation time of 336 min, we collected the sample and heat inactivated the nickase and polymerase for 30 min at 95 °C to stop further exponential amplification of the parasite.

To study the degradation behaviour of the parasite, we relied on the fact that the emerged sequences are tandem[2] and quasi-palindromic repeats[3] of the nicking recognition site. By cleaving the parasite at these repetition sites, one could break down the long parasitic chains into shorter strands. While the nickase would only be partially efficient at this task, since it can only cut one strand of the double-stranded DNA (dsDNA), the BsmI restriction endonuclease can cleave both strands of the recognition site. To prove that parasite can be degraded by the respective restriction endonuclease, we incubated a sample of the parasite created in Figure S1a at 37 °C in the absence or in the presence of BsmI (restriction endonuclease with same binding site as the Nb.BsmI nickase) for 3.5 h (Figure S1b). As predicted, only in the presence of BsmI the EvaGreen (EG) fluorescence (a dsDNA fluorescent marker) significantly decreases across time, implying the cleavage of parasitic dsDNA. Gel electrophoresis analysis confirmed the significant reduction in the presence of long dsDNA parasitic chains, and the apparition of shorter strands that we attributed to bi-products of the cleavage of the longer strands (Figure S1c).

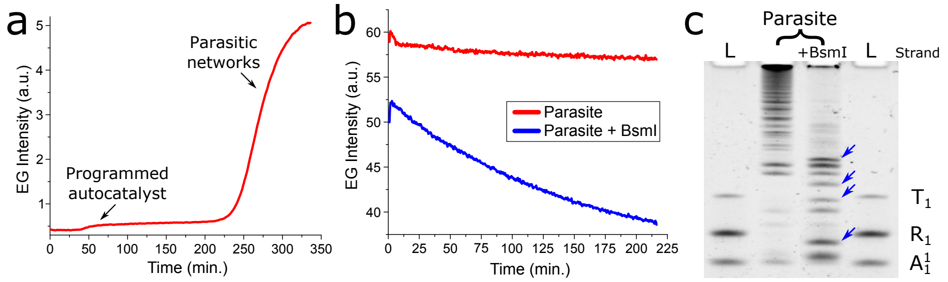


Figure S1: Emerging parasites can be cleaved by restriction endonucleases. (a) EvaGreen fluorescence (dsDNA marker) *versus* time for the incubation of 16 U/mL Bst polymerase, 100 U/mL Nb.BsmI nickase and 50 nM of Template $\mathbf{T}_1$ at 44 °C. (b) EvaGreen (EG) fluorescence *versus* time for samples from panel a incubated in the absence or in the presence of 200 U/ml of BsmI (restriction endonuclease with homologous binding site to Nb.BsmI) at 37 °C. (c) Denaturing polyacrylamide gel electrophoresis (PAGE) showing the multi-band appearance of parasitic networks after its incubation in the absence of the presence of BsmI (panel b). Blue arrows indicate the apparition of shorter strands after BsmI incubation. **L** is a ladder containing template $\mathbf{T}_1$, repressor $\mathbf{R}_1$ and species $\mathbf{A}_1^1$. Conditions panel a: *Kin* buffer with 0.1 mM DTT

Next, we studied the emerging parasitic DNA reactions in the presence of FBS. In particular, we were interested in understanding if the emerging parasitic networks were tandem repeats of the Nb.BsmI nickase (as observed in Figure S1c), if they were developing from nucleases present in the FBS that give rise to parasite (MT Figure 1c), or both simultaneously (multienzymatic). To do so, we collected samples at the end of the experiment of MT Figure 1b for the 10% FBS condition and heat inactivated them to denature the Nb.BsmI and any proteins from the FBS. Next, we incubated the samples at 37 °C either in the absence or in the presence of BsmI, 10% FBS or both (Figure S2a). We observed that the behaviour differed from previous results, where now the incubation with BsmI had little reduction on EvaGreen fluorescence. On the other hand, the addition of 10% FBS drastically reduced the signal, before a sudden increase (which is outside the nature of this manuscript to understand). Gel electrophoresis analysis revealed that the parasite emerged in 10% FBS had a smear form rather than the characteristic multi-band appearance of parasitic networks (Figure S2b), revealing a different nature of parasite. This is further reinforced by the low degradation efficiency when incubated with BsmI and the high degradation by FBS incubation. Nevertheless, we observed that complete parasite degradation only occurred when the parasitic sample was incubated in both BsmI and 10% FBS (Figure S2c), probably indicating the emergence of a multienzymatic parasite requiring both Nb.BsmI and another nuclease present in the FBS.

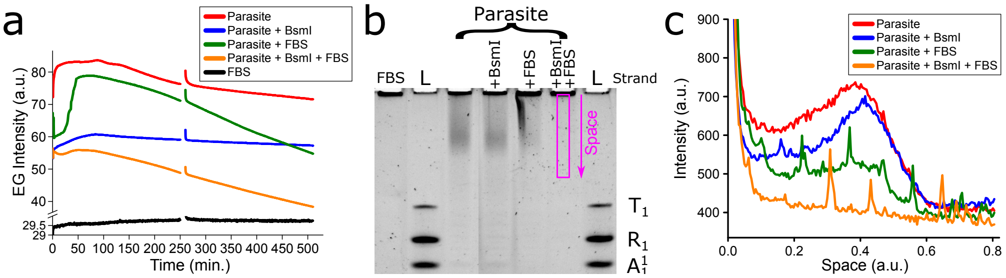


Figure S2: Exponential parasites evolved in 10% FBS differ from traditional parasites evolved in conventional DNA buffers. (a) EvaGreen fluorescence *versus* time for 10% FBS samples from MT Figure 1b incubated in the absence, in the presence of 200 U/ml of BsmI, 10% FBS or both BsmI and 10% FBS at 37 °C. The jump at the 250 min is due to the reset of the thermal cycler. (b) Denaturing PAGE of the incubated samples of panel a. (c) Profiles of the lanes of panel b at the height depicted by the pink rectangle.

### 3 Higher robustness of Nb.BssSI to cellular media

In comparison to the Nb.BsmI network used in MT Figure 1b, we observed that the parasite-resistant Nb.BssSI network (MT Figure 2c) was 3-fold faster in the onset of the exponential amplification and had a 4.5-fold shorter exponential amplification time in the absence of FBS. Although this difference can be attributed to sequence design,[4] we noticed that the enzymatic concentration of polymerase and nickase in Nb.BssSI reactions are 1.25-fold and 5-fold lower, respectively, than for Nb.BsmI reactions. Enzymatic rate experiments revealed that the Nb.BssSI nickase is 36-fold faster than Nb.BsmI in the absence of FBS (Figure S3a and Figure S4), explaining the increase in dynamics of the network even with lower enzymatic concentrations. Upon the addition of 10% FBS, the enzymatic activity of Nb.BssSI decreases by 5-fold, a decrease that explains the delay previously observed in MT Figure 2c, but that is still 15-fold faster than Nb.BsmI.

Taking advantage of the high enzymatic activity of Nb.BssSI, we decided to push further the biocompatibility limits of the actual buffer. First, we assessed the possibility of increasing the concentration of the cell culture rich medium, as it contains the salts and energy source required for sustained *in vitro* cell culture. While previously we had observed a loss in the exponential amplification behaviour with the increase in Dulbecco's modified Eagle's rich medium (DMEM) concentration,[5] the Nb.BssSI network is more robust to chemical perturbations as it stills conserves sigmoidal shape at high DMEM concentrations (Figure S3b). 0.89x DMEM concentration was chosen as it is the standard concentration used in cell culture (when supplemented with 10% FBS and 1% antibiotics). We also noted that this behaviour was still conserved in other standard rich media such as RPMI-1640 (Figure S7). Secondly, we screened for un-essential components that are used to stabilize the PEN DNA toolbox[4] but that could present toxicity to cells, such as for example netropsin that is used to delay the apparition of parasite. Due to the removal of EvaGreen (due to its cytotoxicity as DNA intercalator) from the buffer, we used a yellow fluorophore conjugated to the DNA template to follow the exponential amplification. In this configuration, the quenching of the fluorophore occurs upon template hybridization (Figure S8), and hence the shift in fluorescence signal is related to [dsDNA].[6] Figure S3c shows that the dynamics of the Nb.BssSI network are mainly dependent on the $MgSO_4$ and dithiothreitol (DTT) concentration in a rich medium with 10% FBS solution. We decided to conserve a 6 mM $MgSO_4$ concentration as exponential amplification is drastically affected under 4 mM, and 6 mM is 7-fold greater than that found in DMEM but still half of what has been described for DNA networks in the presence of FBS.[7] We have names this new buffer as *Cell+* buffer due to its higher composition on cell culture medium.

To test the effect of the new *Cell+* buffer on the enzymatic activity, we performed the same enzymatic rate experiments for both nickases in the *Cell+* buffer (Figure S3d). We found that, as in the previous case, the Nb.BssSI nickase is 39-fold faster than the Nb.BsmI. On the other hand, upon the addition of 10% FBS, both enzymatic rates decreased similarly, and as a consequence Nb.BssSI was still 36-fold faster than Nb.BsmI. In addition, we observed that there was a general decrease (between 2 and 5-fold) in enzymatic rates from the previously defined buffer and the new screened buffer. For this reason, to distinguish between both buffers (see table S1), we have termed in this manuscript "*Kin* buffer" to the former buffer due to its higher enzyme kinetics, and "*Cell+* buffer" to the new buffer due to its higher composition in cell culture medium.

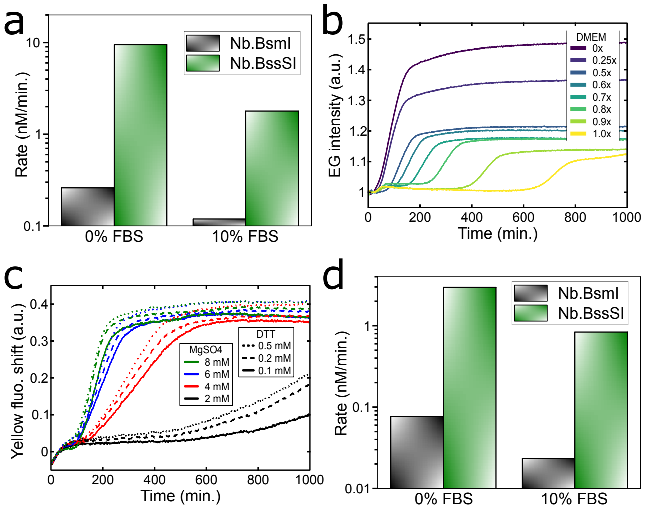


Figure S 3: Measured enzymatic kinetics and buffer screening for higher biocompatible buffers. a) Enzymatic rates for Nb.BssSI and Nb.BsmI in the *Kin* buffer in the absence or the presence of 10% FBS. b) EG fluorescence *versus* time for the exponential amplification of strand $\mathbf{A}_2$ in a concentration range of DMEM. c) Fluorescence shift from the fluorescently-labeled $\mathbf{T}_2$ *versus* time showing PEN reactions are largely dependent on $MgSO_4$ and partially on DTT concentrations in DMEM with 10% FBS. d) Enzymatic rates of the nickases in the new *Cell+* buffer with 0% and 10% FBS. Data in panels a and d determined from degradation experiments with 40 U/ml of nickase at 37 °C (Figure S4). Conditions panel b: *Kin* buffer (in the absence of netropsin) with 1 mM DTT in the presence of 10% FBS.

### 4 Table S1 to S2

Table S1: Composition of the two buffers used in this work. The $X$ in d$X$TPs stands for either N (when dATP, dTTP, dCTP and dGTP) or 3 (when it lacks dATP) nucleotides used. Note that in the presence of 10% FBS, the DMEM in the cell growth medium is reduced down to 0.89x.

| **Name** | **Component** | Concentration |
|---|---|---|
| Cell culture growth medium | | |
| | DMEM | 0.99x |
| | Antibiotics | 1% |
| *Kin* buffer | | |
| | BSA | 0.125 g/l |
| | DMEM | 0.5x |
| | Antibiotics | 1% |
| | $MgSO_4$ | 8 mM |
| | d$X$TPs | 0.8 mM |
| | Syperonic F108 | 0.1% |
| | Netropsin | 2 uM |
| | Tris-HCl | 20 mM |
| | KCl | 10 mM |
| *Cell+* buffer | | |
| | BSA | 0.125 g/l |
| | DMEM | 0.89x |
| | Antibiotics | 1% |
| | $MgSO_4$ | 5.27 mM |
| | d$X$TPs | 0.8 mM |

Table S2: DNA sequences used in this work. Asterisk '*' are phosphorothioate bonds, '**p**' are terminal phosphates, 'JOE' and 'FAM' are fluorophores and 'Dabcyl' is the quencher of FAM. The left column shows the name of the species used in the Main Text, while the right column the name used in the lab. Subscripts define nodes of different networks (based on sequence), while the superscript '1' indicates the addition of a 5' phosphate to the same sequence and the superscript '*' complementarity to the given sequence.

| **Name** | **Sequences 5' → 3'** | **Lab name** |
|---|---|---|
| $\mathbf{A}_1$ | CATTCTGCGAG | Ba-A8 |
| $\mathbf{A}_1^1$ | **p**CATTCTGCGAG | pBa-A8 |
| $\mathbf{T}_1$ | JOE-*C*T*C*GCAGAATGCTCGCAGAA**p** | JOE_CBa-A8(-2)PS3 |
| $\mathbf{R}_1$ | T*T*T*TCTCGCAGAATG**p** | pTBa-A8_T4 |
| $\mathbf{A}_2$ | TCGTGTTCTTC | nA6 |
| $\mathbf{T}_2$ | JOE-*G*A*A*GAA*C*A*CGAGAAGAACAC**p** | JOE_TnA6-2 |
| $\mathbf{R}_2$ | A*A*A*AGAAGAACACGA**p** | pTnA6_A4 |
| $\mathbf{R}_2^*$ | TCGTGTTCTTCTTTT | pTnA6_A4* |
| random | CATCTTCATCCCATCTTCATCC | Lp*Lp* |
| $\text{ref}_1$ | FAM-CCGCATTCGACTCAGAAAAAAAAAACTGAGTCGAATGCGG-Dabcyl | BsmICheck |
| $\text{ref}_2$ | FAM-CGCTCGTGGATCCAGAAAAAAAAAACTGGATCCACGAGCG-Dabcyl | BssSICheck |

### 5 Figures S4 to S23

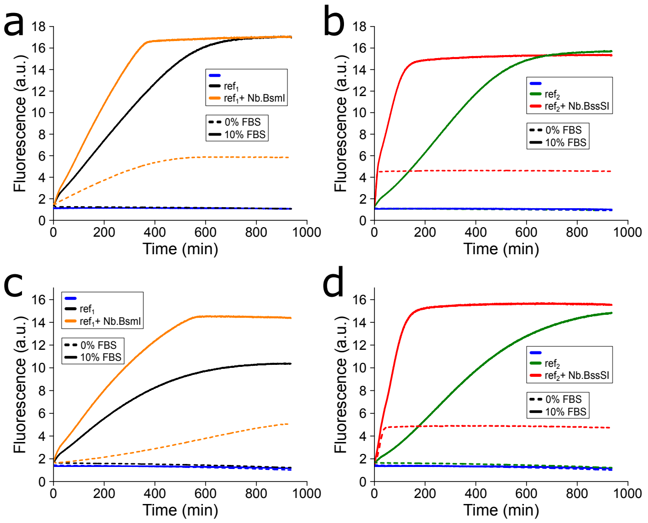


Figure S4: Nb.BssSI conserves higher enzymatic activity than Nb.BsmI in the presence of FBS. Fluorescence *versus* time for the nicking rate assessment in the *Kin* buffer in the absence or presence of 10% FBS for Nb.BsmI (a) and Nb.BssSI (b), and in the *Cell+* buffer for Nb.BsmI (c) and Nb.BssSI (d). Conditions: 40 U/ml of the nickase was incubated with 100 nM of its respective reference substrate at 37 °C.

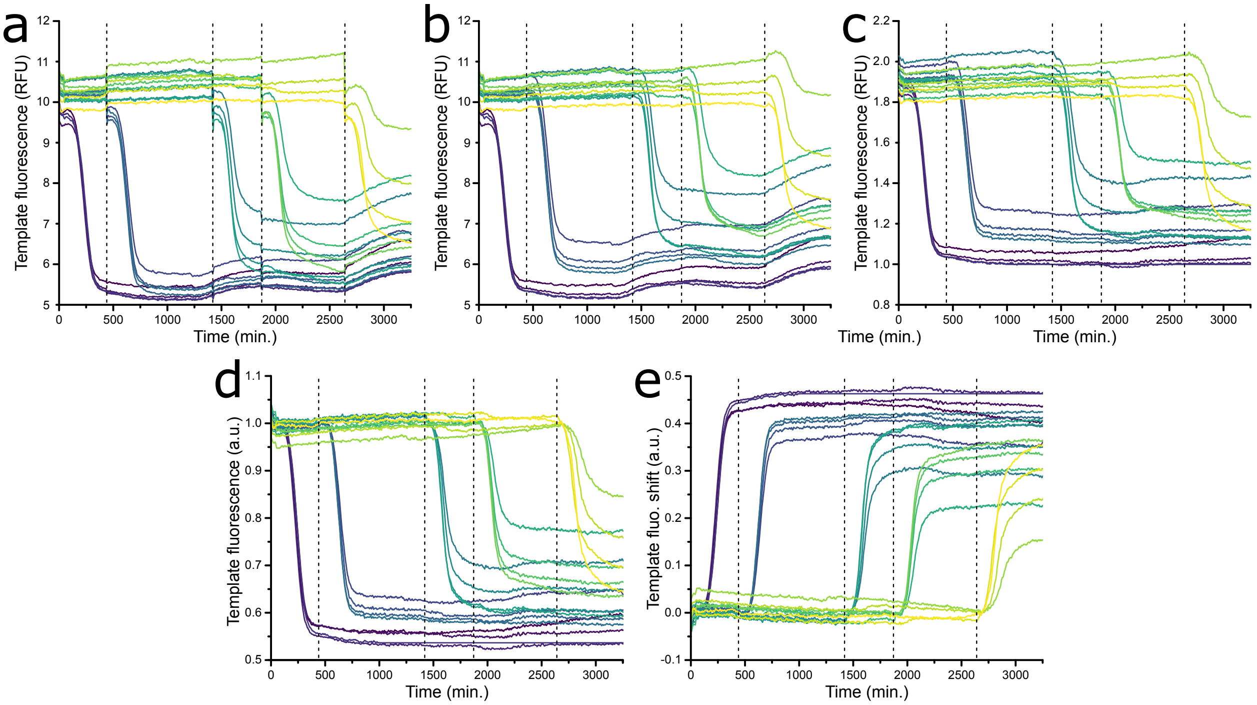


Figure S5: Data treatment used in this work to remove fluorescent artifacts. a) Raw fluorescent data obtained from a Qiagen Rotor-Gene qPCR results for Figure S6. b) Fluorescent data after the fluorescent jumps caused by the opening of the thermal cycler have been removed. c) Results after the removal of the long thermal fluctuations, by dividing the fluorescent data by a reconstructed baseline curve. d) Correction of the fluorescence intensity by an early time point after its activation. e) Final template fluorescence shift values (denoted Yellow fluo. shift in the rest of the figures). Note the straight line of one of the curves after the 1000 minute time point has been kept here for direct comparison with Figure S6. The vertical dashed lines depict the opening of the thermal cycler.

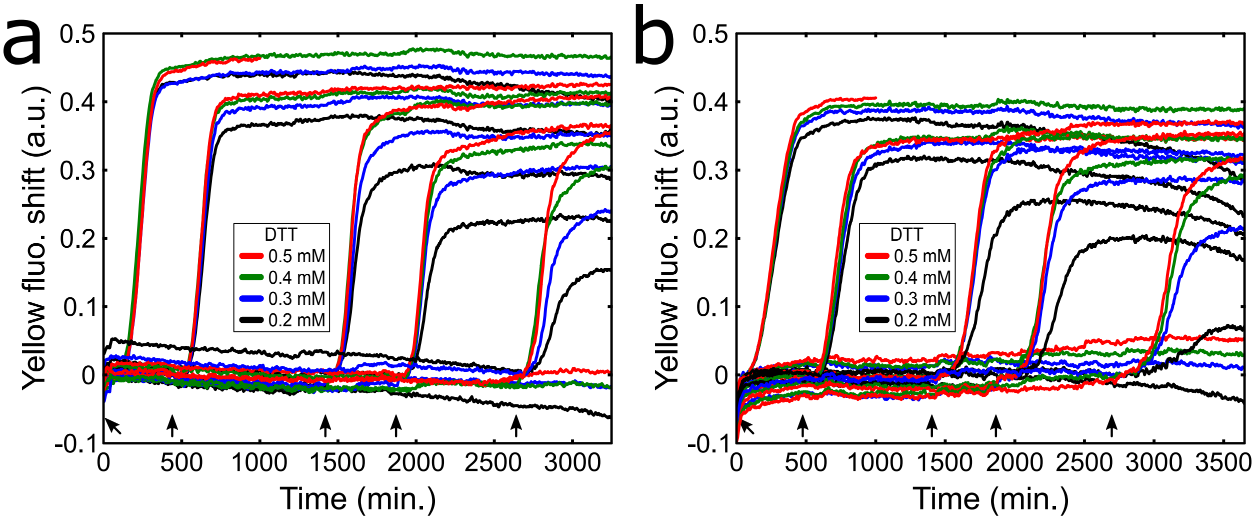


Figure S6: The responsive behaviour of the DNA program in different buffers associated to MT Figures 3 and 4. Fluorescent shift from fluorescently-labelled $\mathbf{T}_2$ *versus* time for the response of the program in a concentration range of DTT in *Kin* buffer (a) and *Cell+* buffer (b). Note that one set of DTT concentration range has not been activated to show the robustness of the *OFF* state. Please refer to SI Section 1.2 for the absence of the curve after 1000 minutes for the 0.5 mM DTT condition activated at t = 0 h. Conditions are identical to MT Figures 3 and 4.

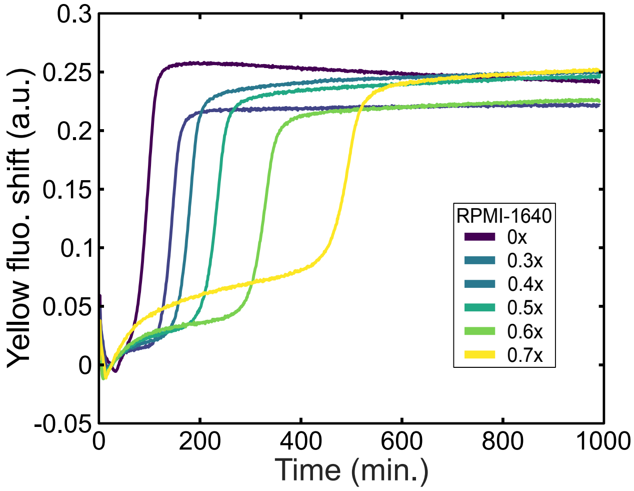


Figure S7: PEN reactions conserve exponential amplification when RPMI-1640 is used instead of DMEM. EG fluorescence *versus* time for the exponential amplification of strand $\mathbf{A}_2$ in a concentration range of RPMI-1640 (PAN Biotech P04-18500). Conditions: *Kin* buffer (with the substitution of DMEM by RPMI-1640), 1 mM DTT and 10% FBS. $[\mathbf{T}_2]_0$= 50 nM.

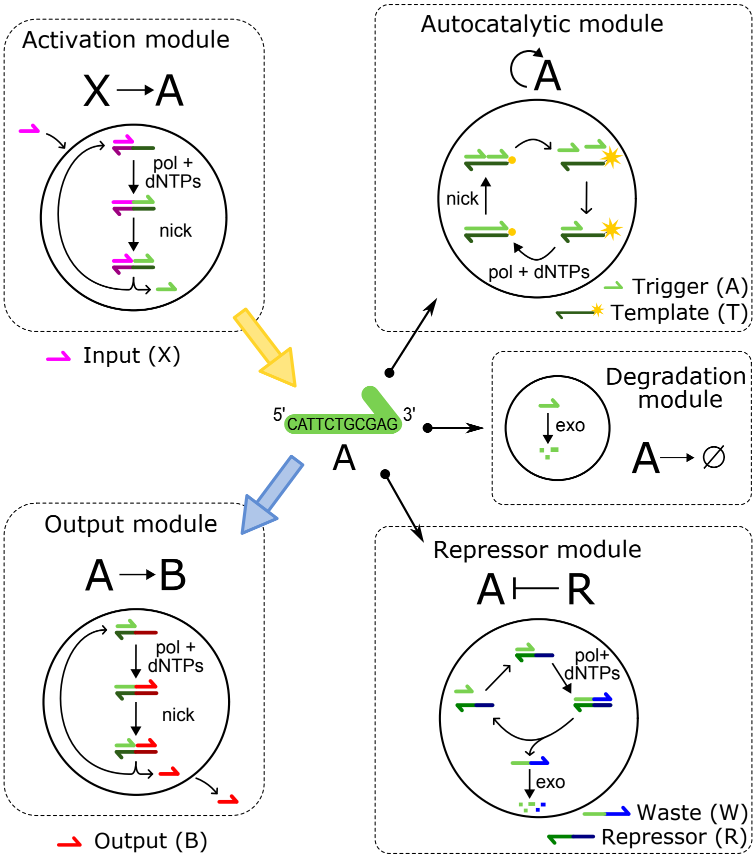


Figure S8: Elementary reactions of the PEN DNA toolbox that can be connected to create functional reaction networks, such as biochemical concentration patterns[8–10] and trigger-driven networks.[11,12] The core mechanism of this DNA program is the exponential amplification of single-stranded DNA (harpoon-ended arrows) by the enzymes polymerase (pol) and nickase (nick), a template (**T**) and the consumption of deoxynucleotides (dNTPs). The enzyme exonuclease (exo) allows the selective degradation of single-stranded DNA, which grants bi-stability behaviour in the presence of a repressor (**R**). The elongation of the trigger (**A**) on the template **T** causes the quenching of the fluorescent marker (yellow star) attached at the 5' of the template.

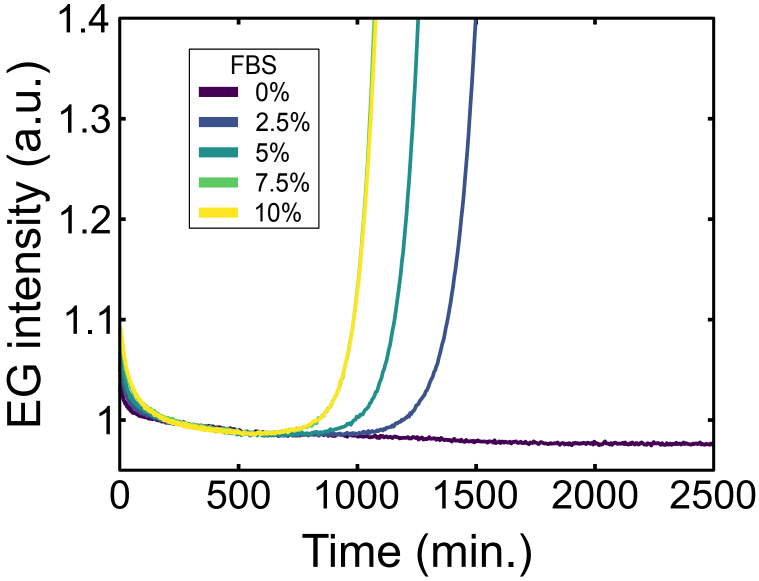


Figure S9: Parasite apparition in the presence of polymerase is enhanced by FBS concentration. Incubation of 8 U/ml Bst polymerase and dNTPs in a gradient of FBS in the *Kin* buffer and 0.1 mM DTT.

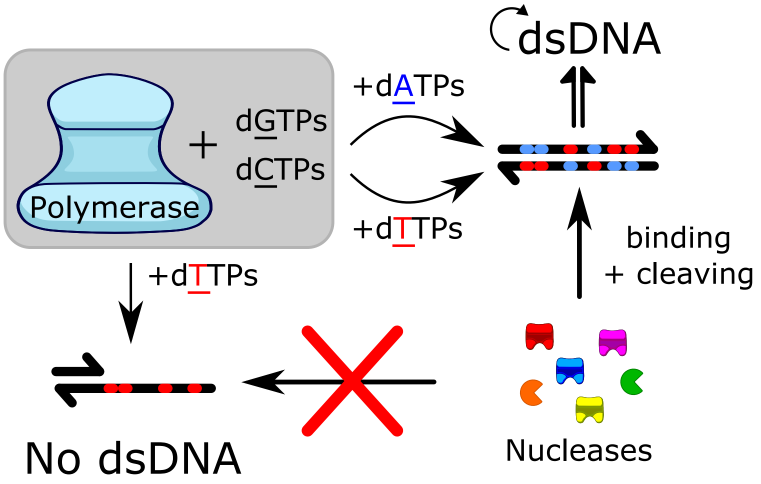


Figure S10: In the presence of polymerase, nucleases and the four standard deoxynucleotides (dATP, dTTP, dCTP and dGTP) autocatalytic amplification of unprogrammed dsDNA (parasites) occurs. This amplification emerges from the *de novo* synthesis of dsDNA by the polymerase and the apparition of recognition sites within these dsDNA, which are cleaved by the nucleases into smaller dsDNA nucleation sites. In the absence of one nucleotide (e.g. dATP) a dsDNA containing the recognition site cannot be created, impeding parasite development.

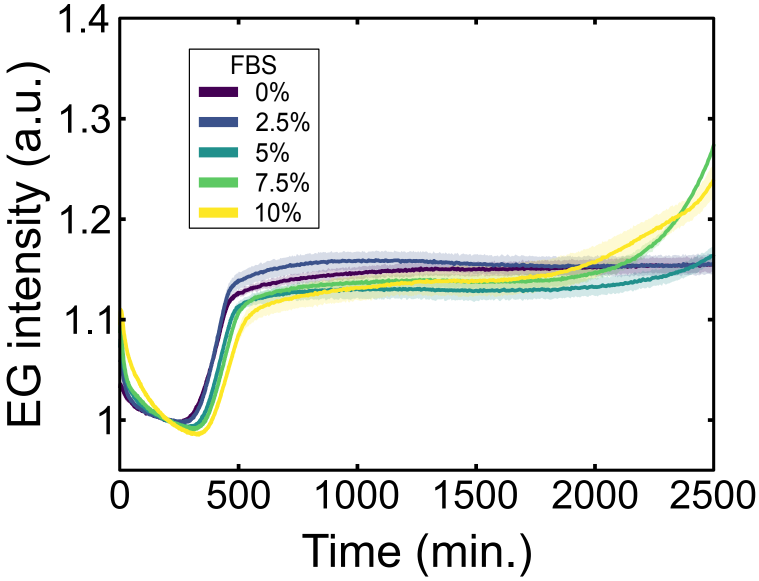


Figure S11: Parasite is also enhanced by FBS in a three-letter code network when using standard dNTPs mix. Conditions are identical to MT Figure 2c but using 4 nucleotides (d<u>A</u>TP, d<u>T</u>TP, d<u>G</u>TP and d<u>C</u>TP).

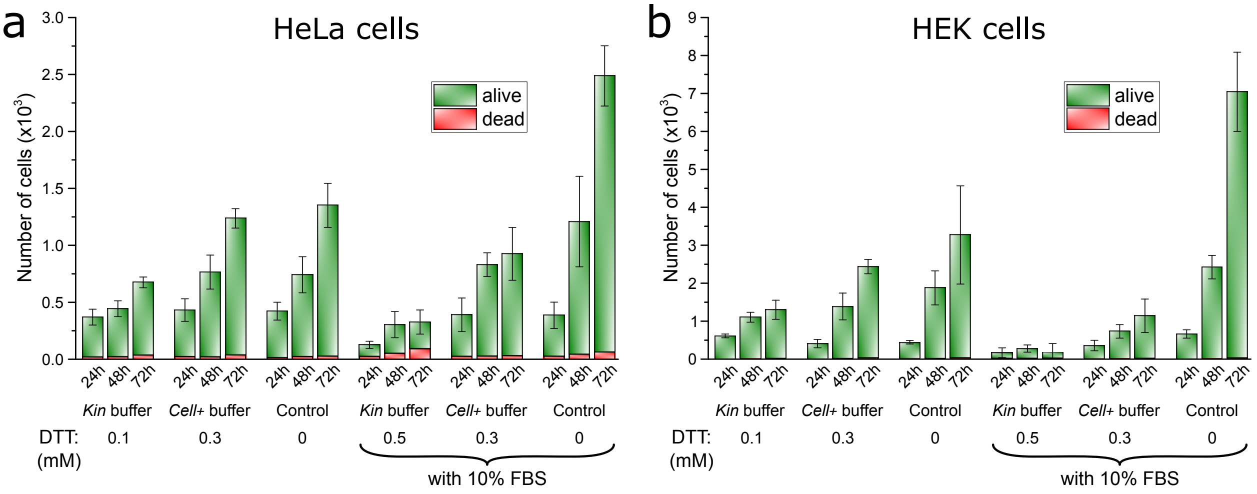


Figure S12: Quantification of living and dead cells determined by propidium iodide staining and flow cytometry for HeLa cells (a) and HEK cells (b) incubated in different buffers and for different incubation periods. Experiments performed per triplicate.

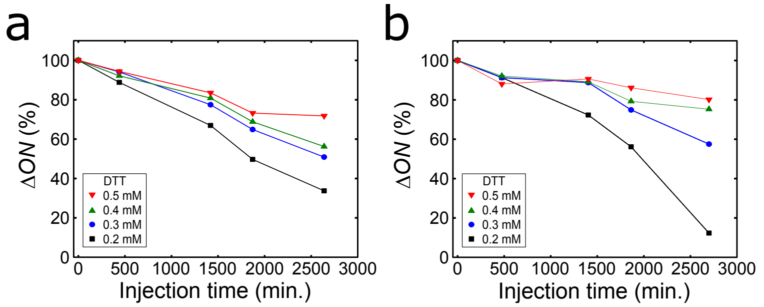


Figure S13: Calculated % of the fluorescence shift of the amplitude of the response ($\Delta I$) of the temporal responsiveness compared to the initial responsiveness at t = 0 h *versus* injection time for the *Kin* buffer (a) and *Cell+* buffer (b). Data determined from Figure S6. Solid lines are guides to the eye.

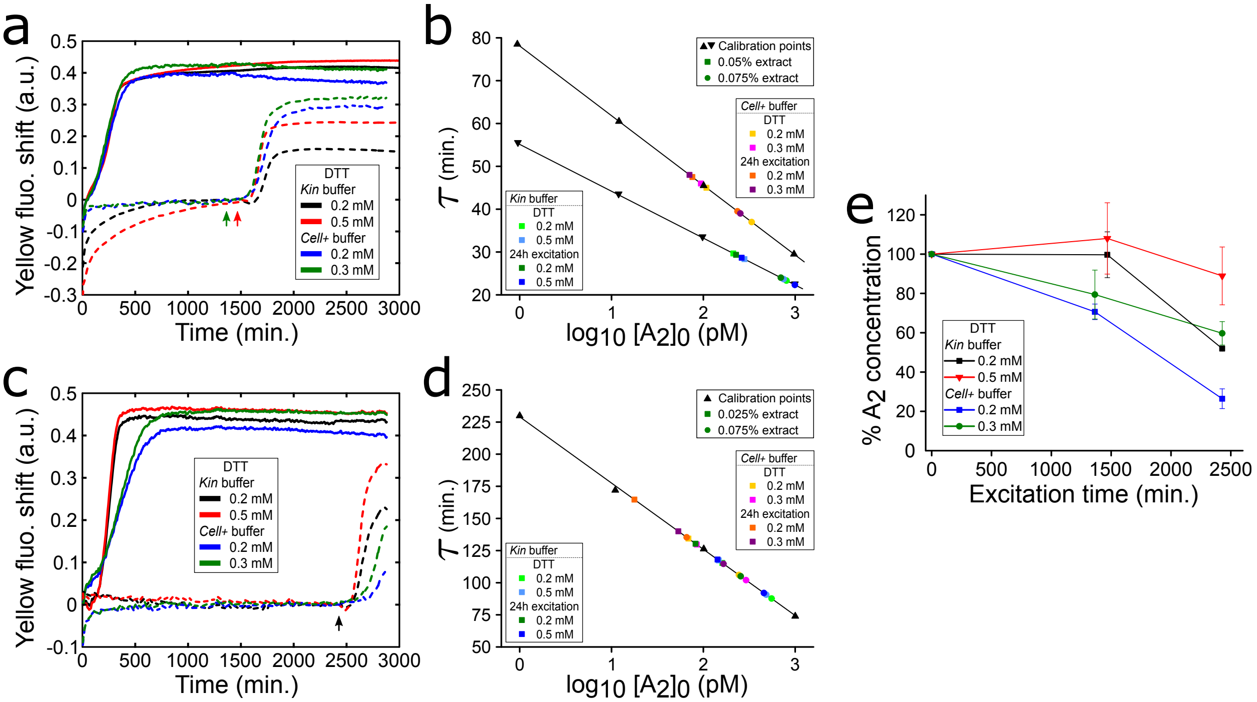


Figure S14: Isothermal quantification of trigger $\mathbf{A}_2$ production at $\sim$49 h after different activation time points. a) Fluorescence shift *versus* time for the $\mathbf{A}_2$ *OFF* state activated at 0 h (solid lines) or after 24 h (dash lines) for *Kin* buffer and *Cell+* buffer with two different DTT concentrations. Experiments were performed on different days, reason why there is a temporal difference between the activation of *Kin* buffer (red arrowhead) and *Cell+* buffer (green arrowhead). b) The amplification onset times ($\tau$) *versus* $\log_{10}[\mathbf{A}_2]_0$ for a trigger titration calibration curve (black triangles) and from samples extracted from panel a and diluted down to 0.025% or 0.075% (square and circle symbols, respectively) into a fresh isothermal amplification. The extracted samples were plotted within a linear fit of the upward pointing triangles and downward pointing triangles for the *Cell+* buffer and *Kin* buffer conditions, respectively, to quantify trigger $\mathbf{A}_2$ concentration. c) *idem* to panel a but for the activation after 40 h (black arrowhead). d) *idem* to panel b but for samples extracted from panel c. e) % trigger $\mathbf{A}_2$ concentration with respect to activation at t = 0 h for the 24 h responsiveness and 40 h responsiveness quantified from panel b and d respectively. All experiment performed at 37 °C. Panel a and c experimental conditions are identical to MT Figures 3 and 4, respectively. We attribute the $\sim$3.5-fold slower $\tau$ in panel d to the presence of 2 uM of netropsin in the isothermal amplification, which causes a reduction in exponential dynamics[13] but would not affect trigger quantification as trigger calibration is done in the same condition. Solid lines in panels b and d are linear fits, while in panel e are guides to the eye. Error bars in panel e correspond to the standard deviation of a triplicate experiment.

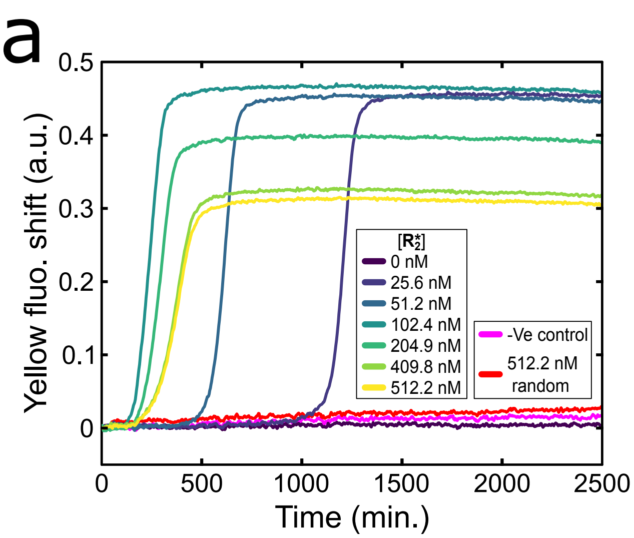


Figure S15: The activation of the DNA program responds non-linearly to the $\mathbf{R}_2^*$ concentration. a) Fluorescent shift *versus* time for the response of 200 nM $\mathbf{T}_2$ with 100 nM $\mathbf{R}_2$ in *Kin* buffer with 10% FBS and 1 mM DTT, with increasing concentration of $\mathbf{R}_2^*$ and with 512.2 nM of a random sequence (red). -Ve control (pink) is in the absence of polymerase. b) Amplification onset time ($\tau$) after the activation spike at t = 33 minutes *versus* $\mathbf{R}_2^*$ concentration. Non-linear behaviour correlates with the non-linear nature of the repression mechanism,[14] where fastest activation is accomplished at equimolar concentration of $\mathbf{R}_2^*$ to $\mathbf{R}_2$.

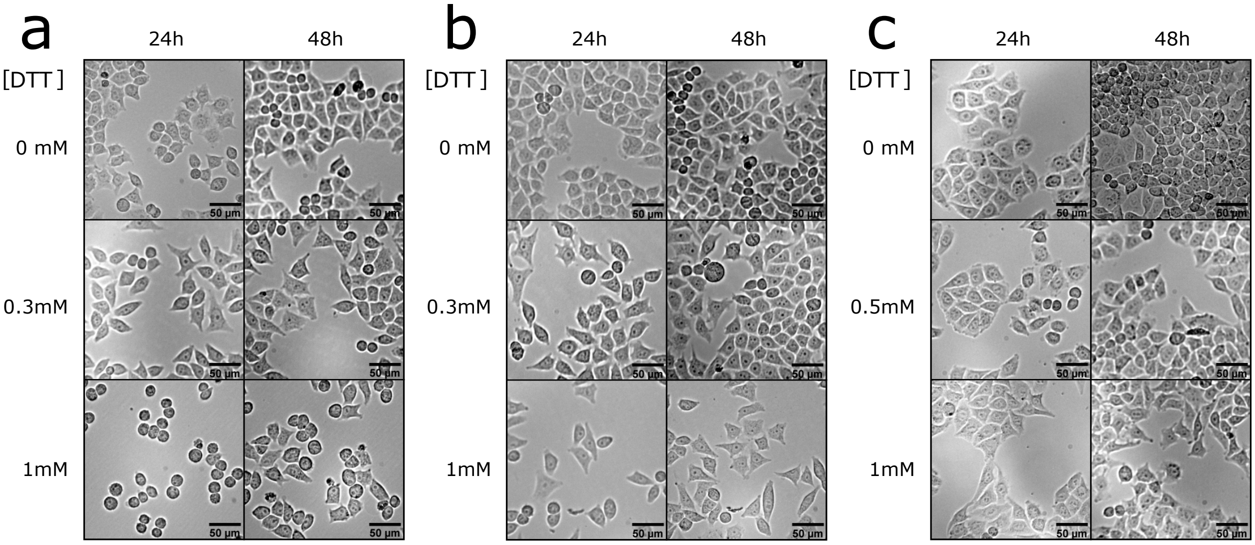


Figure S16: FBS attenuates the adverse effect of DTT on HeLa attachment. Bright-field images showing cellular morphology at different DTT concentrations in the presence of 2.5% (a), 5% (b) and 10% (c) FBS in the cell culture growth medium.

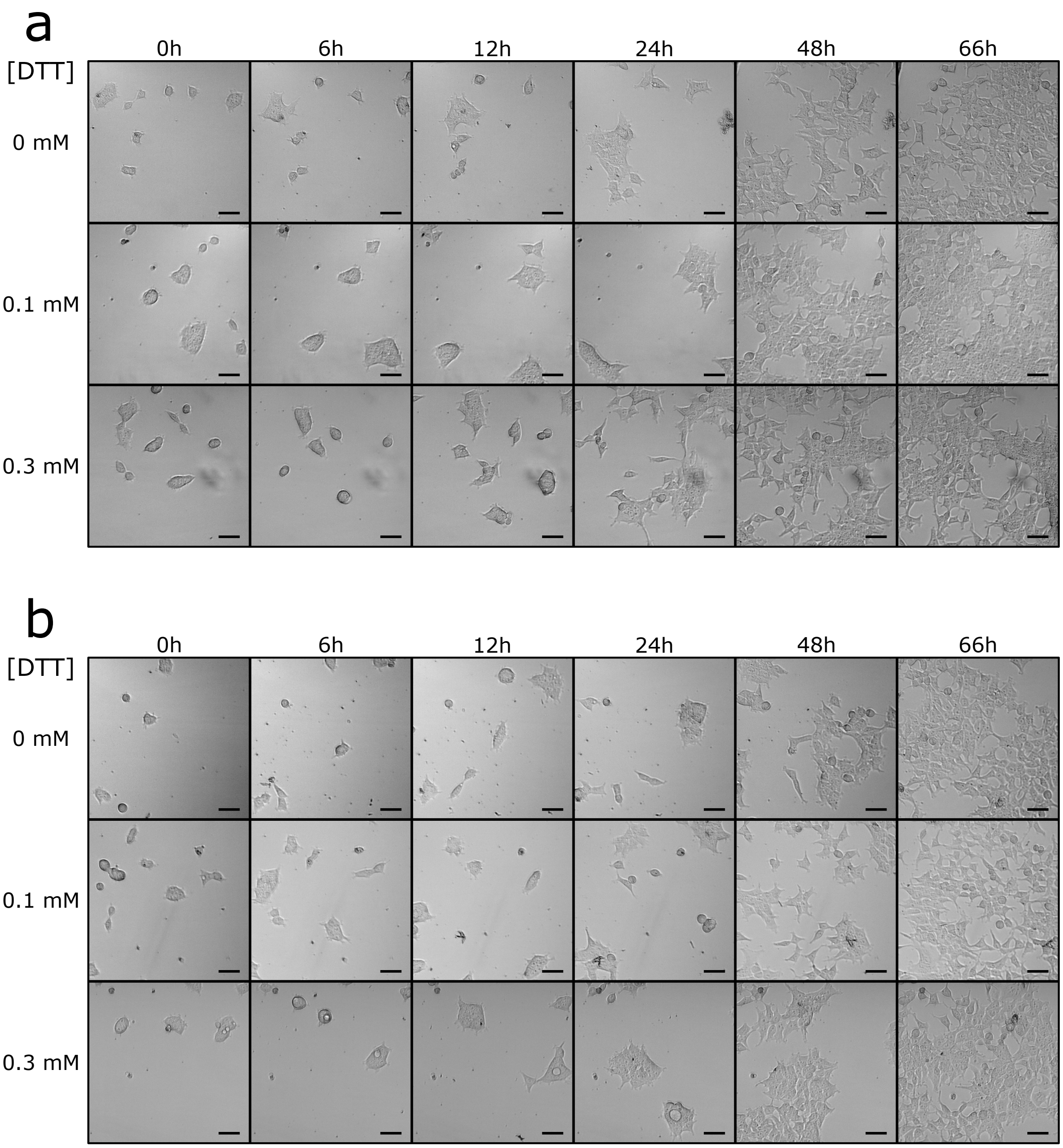


Figure S17: HEK cells still conserve adherent phenotype with 2.5% (a) and 5% (b) FBS in the cell culture growth medium.

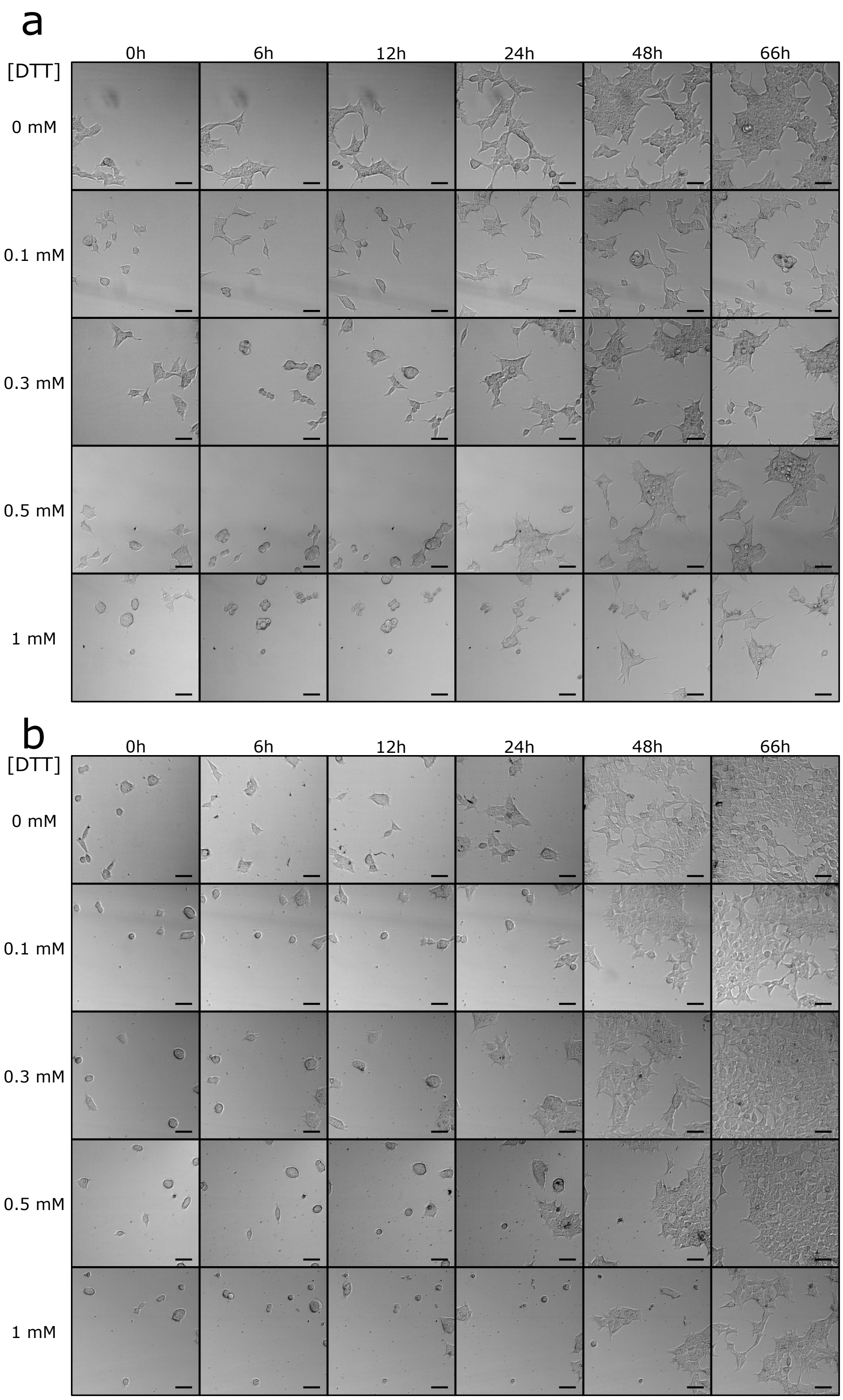


Figure S18: DTT impairs HEK cell growth. Bright-field images showing the aggressiveness of DTT on HEK cells in the absence of FBS (a) and restoration of phenotype (cell extension) in the presence of 10% FBS (b). Experiments performed in the cell culture growth medium.

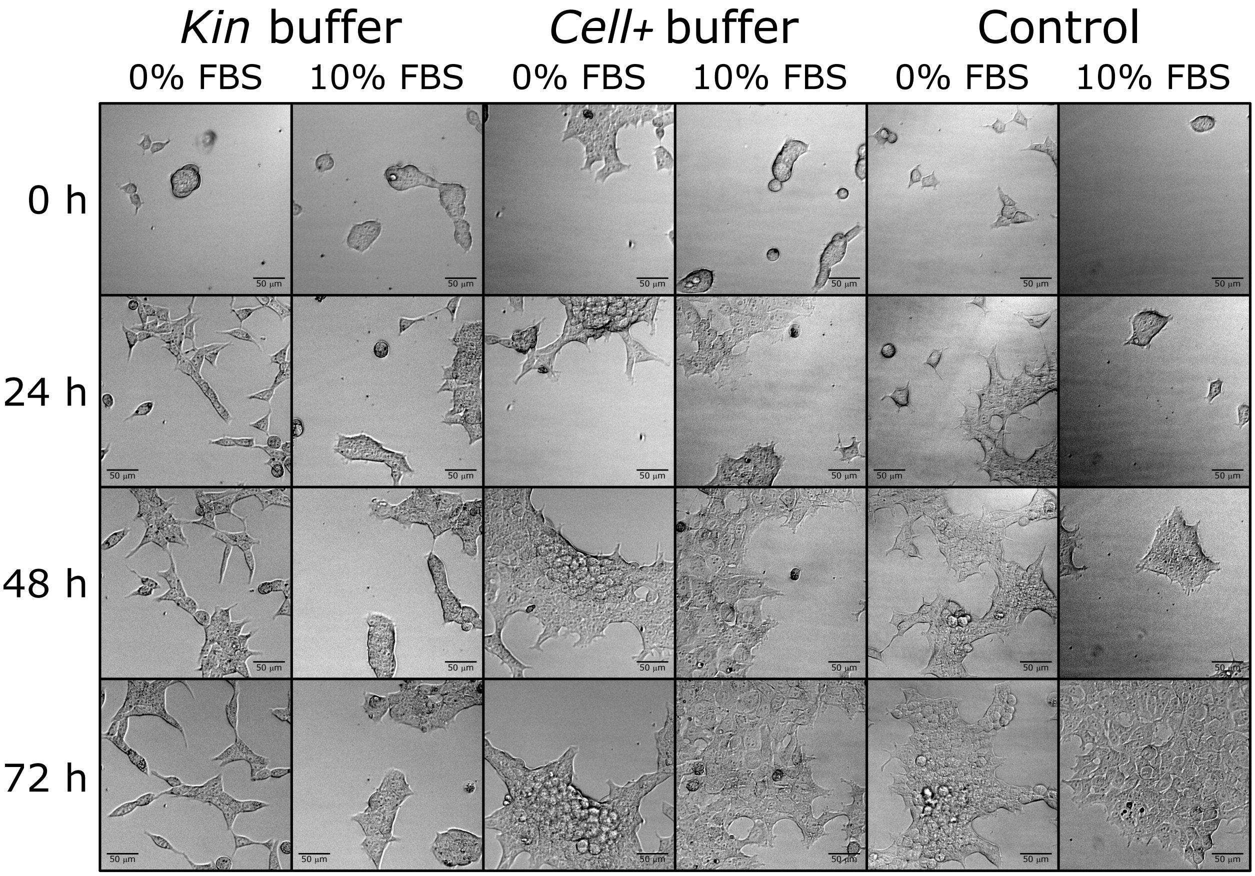


Figure S19: Bright-field images showing the aggregated or stretched phenotype of HEK cells when cultured in the absence or the presence of 10% FBS in the *Kin* buffer, *Cell+* buffer and cell culture growth medium (control).

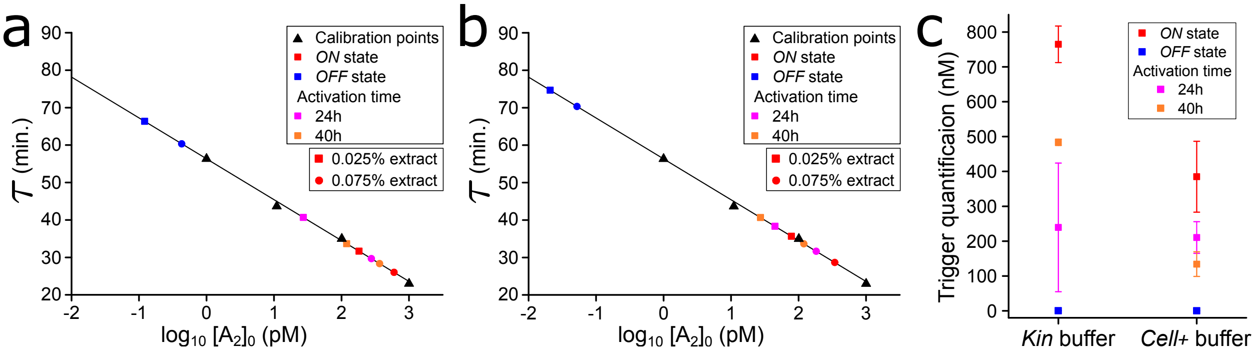


Figure S20: Isothermal quantification of the trigger $\mathbf{A}_2$ present at the end of the experiment of MT Figure 6 and Figure S21a. Samples were extracted from each condition and diluted down to 0.025% or 0.075% (square and circle symbols respectively) into a fresh isothermal amplification. The amplification onset times ($\tau$) were plotted within a trigger titration calibration curve for *Kin* buffer (a) and *Cell+* buffer (b). Panel c shows the predicted $\mathbf{A}_2$ trigger concentrations after 71h of incubation of the DNA/enzyme-based molecular program in the presence of cells and 10% FBS. As obtained in the absence of cells (Figure S14), the $\mathbf{T}_2$ autocatalytic network conserves higher DNA production in *Kin* buffer than in *Cell+* buffer in the presence of cells. In addition, we observed that the *in situ* production of $\mathbf{A}_2$ DNA after activation at 24 and 40 h is ~2-fold lower than the *ON* state condition in both buffers. Analysis of the *OFF* state conditions revealed concentrations under 0.6 nM, ~3 orders of magnitude smaller than the *ON* state. Error bars in panel c are calculated from the average of the 0.025% and 0.075% dilutions.

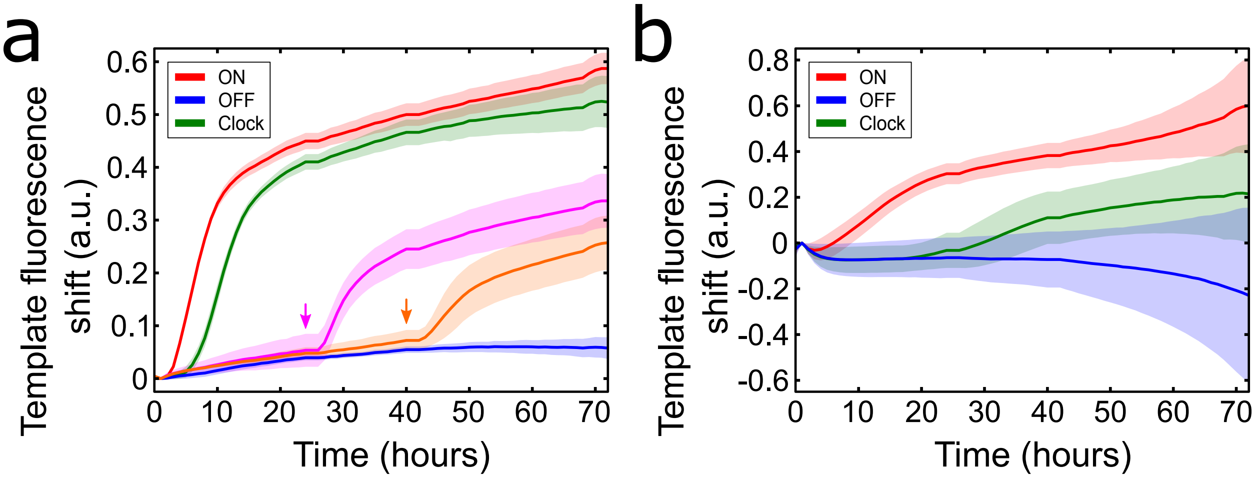


Figure S21: Responsive and pre-programmable DNA programs in the presence of living cells. a) Fluorescence shift of $\mathbf{T}_2$ *versus* time showing the production dynamics of $\mathbf{A}_2$ in the *Kin* buffer with 0.5 mM DTT and in the presence of 10% FBS and HeLa cells. The curves show the unsuppressed *ON* state (red), the pre-programmed clock reaction (green) and the repressed OFF state (blue) and its responsiveness by the addition of $\mathbf{R}_2^*$ at 24 h (pink) and 40 h (orange). Arrowheads indicate the addition time of DNA activator $\mathbf{R}_2^*$. b) Fluorescence shift of $\mathbf{T}_2$ *versus* time for the pre-programmed clock reaction in the *Cell+* buffer with 0.3 mM DTT. *ON* and *OFF* states curves are from MT Figure 6b. The onset of the exponential amplification occurred 3-fold faster in the *Kin* buffer than in the *Cell+* buffer, difference we account to the faster dynamics of the nickase (Figure S4), which causes the autocatalytic reaction to go faster (Figure S22) and hence more robust to repressor (Figure S23). Conditions: The *ON* and *OFF* states started with $[\mathbf{R}_2]_0 = 0$ nM and 150 nM, respectively, while the clock reaction with $[\mathbf{R}_2]_0 = 50$ nM. The DNA activator $\mathbf{R}_2^*$ was introduced at 300 nM.

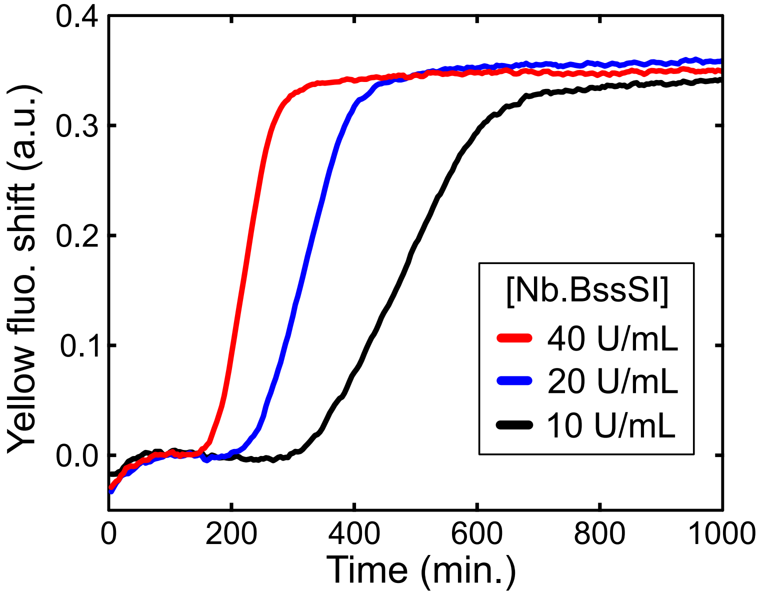


Figure S22: $\mathbf{A}_2$ autocatalytic dynamics are dependent on Nb.BssSI concentration. Conditions: 200 nM $\mathbf{T}_2$ in the *Kin* buffer with 0.5 mM DTT and 10% FBS.

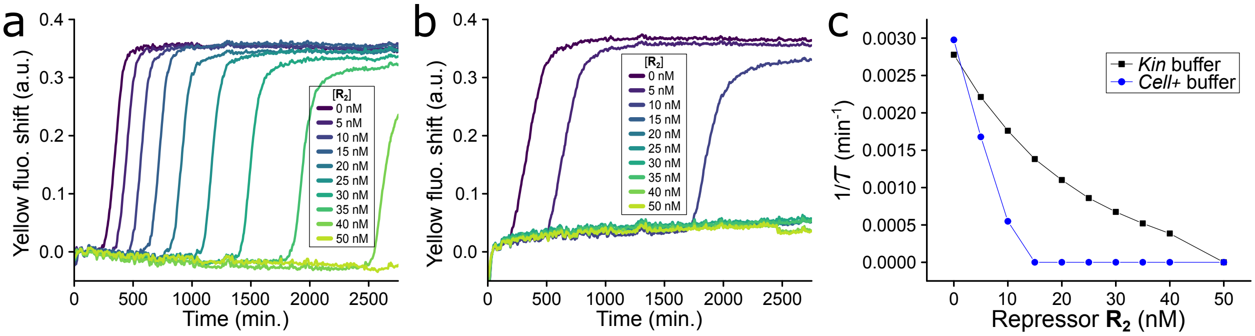


Figure S23: Bistability dynamics of network $\mathbf{A}_2$ in the *Kin* buffer and *Cell+* buffer with 10% FBS. Fluorescent shift *versus* time for the $\mathbf{A}_2$ autocatalyst in a range of repressor $\mathbf{R}_2$ and 10% FBS either in the *Kin* buffer (a) or the *Cell+* buffer (b). c) $1/\tau$ *versus* repressor $[\mathbf{R}_2]$ for the *Kin* buffer (black) and *Cell+* buffer (blue) determined from panel a and b. Solid lines in panel c are guides to the eye.
